# Coevolutionary analyses require phylogenetically deep alignments and better null models to accurately detect inter-protein contacts within and between species

**DOI:** 10.1101/014902

**Authors:** Aram Avila-Herrera, Katherine S. Pollard

**Author notes:** < >.

## Abstract

When biomolecules physically interact, natural selection operates on them jointly. Contacting positions in protein and RNA structures exhibit correlated patterns of sequence evolution due to constraints imposed by the interaction, and molecular arms races can develop between interacting proteins in pathogens and their hosts. To evaluate how well methods developed to detect coevolving residues within proteins can be adapted for cross-species, inter-protein analysis, we used statistical criteria to quantify the performance of these methods in detecting inter-protein residues within 8 angstroms of each other in the co-crystal structures of 33 bacterial protein interactions. We also evaluated their performance for detecting known residues at the interface of a host-virus protein complex with a partially solved structure. Our quantitative benchmarking showed that all coevolutionary methods clearly benefit from alignments with many sequences. Methods that aim to detect direct correlations generally outperform other approaches. However, faster mutual information based methods are occasionally competitive in small alignments and with relaxed false positive rates. All commonly used null distributions are anti-conservative and have high false positive rates in some scenarios, although the empirical distribution of scores performs reasonably well with deep alignments. We conclude that coevolutionary analysis of cross-species protein interactions holds great promise but requires sequencing many more species pairs.

## Background

Coevolution—“the change of a biological object triggered by the change of a related object” [1]—is a powerful concept when applied to molecular sequence analysis because it reveals positional relationships that are worth preserving across evolutionary time scales. Sequence evolution is constrained by essential molecular interactions, such as contacts within a protein or RNA structure, as well as inter-molecular interactions in protein complexes and signaling pathways. These constraints define an epistasis between sites (residues or base-pairs) where the probability of a substitution depends on the states of other sites [2] involved in an interaction. Because epistasis can induce correlation between substitution patterns across columns in multiple sequence alignments, many methods have been developed that use evidence of coevolving alignment columns to detect physical interactions within and between biomolecules. These methods draw inspiration from diverse techniques in molecular phylogenetics, inverse statistical mechanics, Bayesian graphical modeling, information theory, sparse inference, and spectral theory (reviewed in [3, 4]).

Despite good rationale for coevolutionary approaches, physically interacting alignment columns have been notoriously difficult to identify from correlated patterns of sequence evolution for several reasons. First, shared evolutionary history creates a background of correlated substitution patterns against which it can be difficult to distinguish additional constraints derived from physical interactions. Common phylogeny is particularly strong within a gene family (e.g., predicting intra-molecular contacts). But it is also present across gene families within a species or even between species (e.g., predicting host-virus protein interactions), especially at shorter evolutionary distances where gene trees mirror species trees more closely. Coevolution methods have used a variety of approaches to counter the dependence induced by shared phylogeny, including removing closely related sequences from alignments to reduce non-independence [5, 6], differential weighting of sequences when computing statistics [7–9], and null distributions that directly model or indirectly account for phylogeny [10–13].

A second challenge arises when trying to distinguish correlated evolution that arises from direct versus indirect interactions. Alignment columns that are indirectly implicated in an interaction can be strongly correlated, and most columns are involved in multiple, partially overlapping interactions. For these reasons, close physical interactions may not produce patterns of substitution that are significantly more highly correlated than the background present in structures. This problem has been the focus of a recent class of coevolutionary methods that focuses on reducing the number of incorrect predictions by disentangling direct from indirect correlations [9, 14–17]. An alternative point of view considers these networks of indirectly correlated residues as protein sectors that can easily, through cooperative substitutions, respond to fluctuating evolutionary pressures [18].

Finally, due to low power—resulting in part from the previous two challenges—physically interacting sites can typically only be detected in multiple sequence alignments that span large evolutionary divergences and contain many hundreds to thousands of sequences. Recent evaluations of a number of coevolution methods concluded that accurate contact predictions require alignments with one to five times as many sequences (with <90 % sequence redundancy) as positions [19, 20].

To date, coevolutionary prediction of physically interacting alignment columns has been applied with success to intra-molecular contacts [7, 21–23] and well-characterized inter-molecular interactions [24], such as bacterial two-component signaling systems [25], enzyme complexes [26], and fertilization proteins [27]. The signal-to-noise ratio is too low and the search space too large to use sequence evolution to effectively identify pairs of physically interacting protein residues across entire proteomes; most pairs of sites with correlated substitution patterns are not in direct contact, and most physically interacting sites do not have statistically correlated substitution patterns [28].

However, the ability to now measure physical interactions between biomolecules with high-throughput technologies, such as affinity purification followed by mass spectrometry (APMS) [29], two-hybrid methods [30, 31], and protein complementation assays [32], raises the possibility of using sequence coevolution in a more specific way: to refine predicted interactions in an experimentally reduced search space. For example, correlated substitution patterns in pairs of proteins could help determine if an experimentally measured interaction is likely to represent direct physical contact versus an indirect interaction in a complex or a false positive. Coevolutionary analysis could also be informative regarding which of the sites in a pair of interacting molecules are most likely to be in physical contact.

One particularly exciting application of this approach is to characterize and potentially manipulate interacting residues in host-virus and host-parasite protein interactomes [33, 34]. Newly emerging data on antibody and antigen sequences within a host [35] offers an opportunity to harness coevolutionary signals to investigate the mechanisms of broadly neutralizing antibodies and immune evasion. The primary open question for these new applications is whether existing methods are sensitive and specific enough to detect coevolution with the levels of constraint and divergence that are present in inter-molecular data sets of modest size.

To this end, we designed data processing scripts, statistical evaluation and visualization tools, and simulation pipelines that allowed us to easily extend a suite of coevolution methods designed for intra-protein interaction prediction (Table 1) so that they can be used to test for patterns of correlated sequence evolution at pairs of sites in two different proteins, potentially from different sets of organisms in different parts of the tree of life (e.g., human-bacteria, bacteria-phage interactions). We then applied this integrated framework for coevolutionary analysis to refine and annotate a recently derived human-HIV1 protein-protein interaction network [33] and to test for coevolution in the well studied arms-race interaction between the mammalian cytidine deaminase APOBEC3G (A3G) and its HIV1 antagonist, Vif. Because fewer than ten orthologous mammal-lentivirus proteome pairs have been sequenced and mammalian divergence is low, we hypothesized that power would be low in these settings.

**Table 1:**
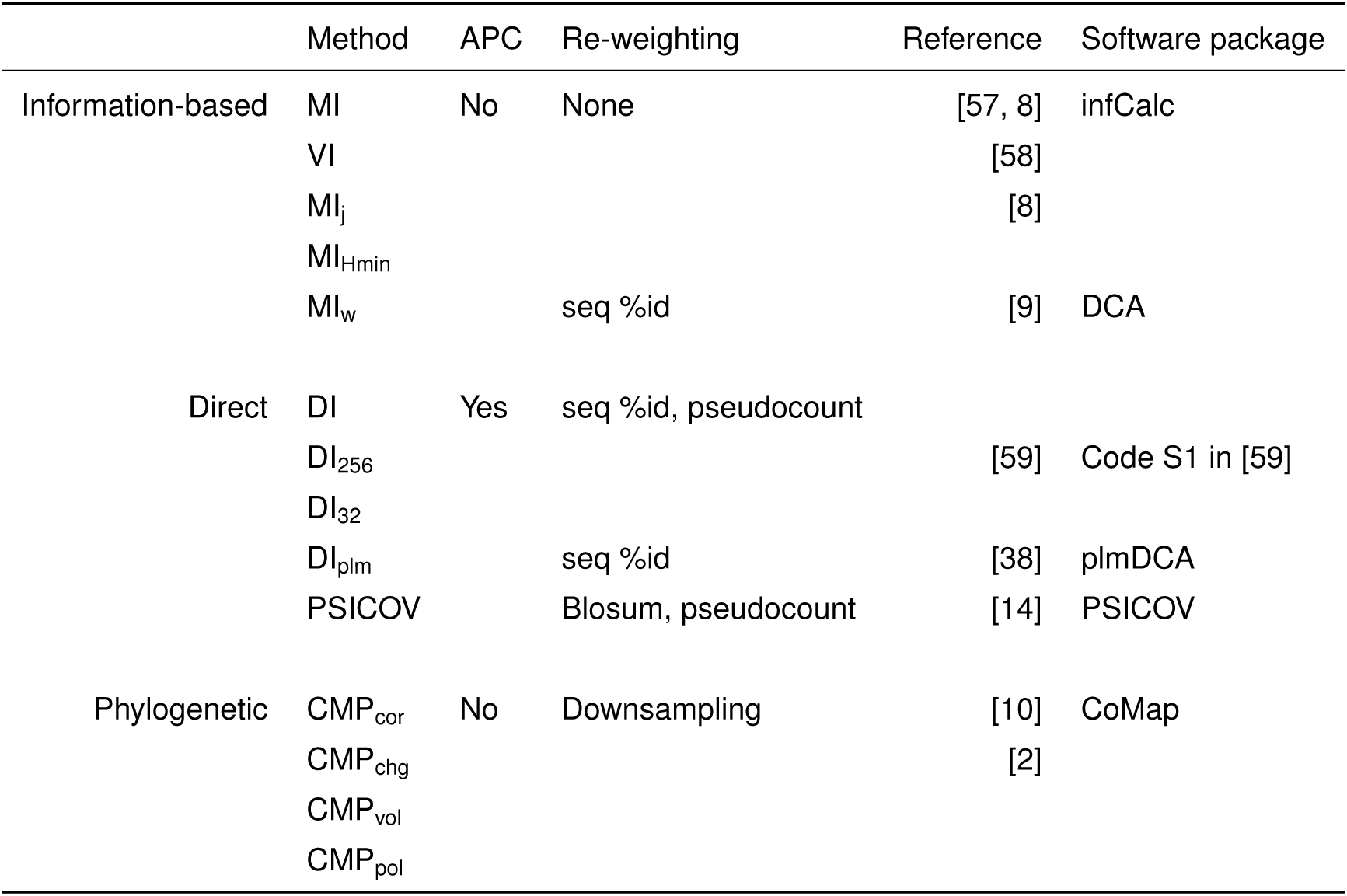
Coevolution methods included in analysis. Information-based methods: MI: mutual information [57], VI: variation of information [58], MI_j_: MI divided by alignment column-pair entropy, MI_Hmin_: MI divided by minimum column entropy [8], MI_w_: MI with adjusted amino acid probabilities. Direct methods: DI: direct information—MI with re-estimated joint probabilities [9], DI_256_, DI_32_: DI using Hopfield-Potts for dimensional reduction (256 and 32 patterns respectively) [59], DI_plm:_ Frobenius norm of coupling matrices in 21-state Potts model using pseudolikelihood maximization [38], PSICOV: sparse inverse covariance estimation [14]. Phylogenetic methods: CoMap *P*-values for four analyses CMP_cor_: substitution correlation analysis [10], CMP_pol_ for polarity compensation, CMP_chg_ for charge compensation, CMP_vol_ for volume compensation [2].

To quantify the limitations of coevolutionary methods when only a handful of sequences are available, we used a data set of 33 within-species bacterial protein-protein interactions. To systematically determine the parameters that affect performance, we focused on the well-characterized interaction between bacterial histidine kinase A (HisKA) and its response regulator (RR), for which a co-crystal structure and thousands of sequences are available. By subsampling HisKA-RR sequence pairs, we show that most methods have appreciable precision or power at low false positive rates for alignments with ~500 or more sequences. However, the best performing method depends on whether power or precision is more important, the number of non-redundant sequences in the alignment, and whether the goal is to find structurally or functionally linked residues. By expanding this analysis to 32 additional bacterial interactions [24], we showed that these trends generalize beyond the specific example of HiskA and RR. We conclude that coevolution methods are able to identify some residues important for cross-species protein-protein interactions, but this approach will benefit greatly from additional sequence data.

## Results

### Performance benchmarking of coevolution methods

The coevolutionary methods benchmarked in our analyses fall into three general groups (Table 1). Information-based methods are various flavors of Mutual Information between pairs of sites, each considered independently. Direct methods are those that consider pairs of sites in the context of a sparse global statistical model for contacts in the multiple sequence alignment. Phylogenetic methods explicitly use a substitution rate matrix and phylogenetic tree in their calculation of a coevolution statistic that may take into account the biochemical and physical properties of amino acid residues, as well as report a *P*-value based on internal simulation of independently evolving sites. In this benchmark we use the CoMap *P*-value as a statistic for comparison with other coevolution methods. Other differences among the coevolution methods include the incorporation of two additional techniques that have been shown to improve performance, re-weighting sequences such that similar sequences contribute less to the final score [5] and applying an Average Product Correction (APC) to remove background noise and phylogenetic signal from “raw” coevolution statistics [8].

To benchmark coevolution methods, we used 33 within-species pairs of proteins with co-crystal structures determined from *E. coli* proteins. These include a set of paired alignments compiled by [24], plus the histidine kinase-response regulator (HisKA-RR) bacterial two-component system from [36], provided by the authors. We included HisKA-RR, because it is a well-characterized interaction with a very large, diverse multiple sequence alignment (8998 sequences for each gene) and genetic evidence supporting several interactions. For these reasons, HisKA-RR has also been used previously in coevolutionary analyses [37].

Because the HisKA-RR alignment is so large, it enabled us to quantify the effects of alignment size and diversity by down-sampling the full alignment to produce a wide range of smaller pairs of HisKA and RR multiple sequence alignments with different numbers of sequences (range 5 to 5000 sequences) and phylogenies from the original alignment. The 32 alignment pairs from [24] naturally varied in size (range 168 to 1428 sequences).

For each pair of multiple sequence alignments from two interacting proteins, we compared every site in the first protein to every site in the second protein and scored these pairs of alignment columns for coevolution using each of the methods in Table 1. We then used coevolution scores to predict inter-domain pairs of amino acid residues that are less than 8 angstroms (Å) to each other, measured between C_β_s, in the representative co-crystal structure (see Methods). We also repeated our analyses of the HisKA-RR sub-alignments using a stricter definition of contacts that requires additional biochemical evidence for specificity determination, and an alternate definition that measures distance between the closest non-hydrogen atoms. Trends in our results were generally similar across these choices of definition for true interactions, but we observed some differences in performance between definitions when enforcing a false positive rate (FPR) (Figure S2).

The performance of each method to distinguish contacting pairs of residues (positives) from other residue pairs (negatives) was measured as previously described [14, 38] and evaluated using power (also called recall and true positive rate (TPR)) and precision (also called positive predictive value (PPV)) at a range of low FPRs. Power and precision are complementary performance measures that quantify the percentage of interacting residue pairs that are found and the percentage of identified residue pairs that are interacting, respectively. Precision is a useful measure of performance in cases where positives (contacting pairs of residues) are overwhelmed by negatives (non-contacting residues). A method with high precision is helpful for generating lists of high confidence pairs of residues for expensive follow-up studies, even if it misses a number of truly interacting sites and therefore has relatively low power. We additionally examined four threshold-independent performance measures, area under Receiver-Operator Curve (auROC), area under precision-recall curve (auPR), maximum F_1_-score (f_max_), maximum *ϕ* (phi_max_).

### Physically interacting sites can be accurately detected in large sequence alignments

Our primary finding is that many coevolutionary methods are able to detect inter-molecular contacts at low FPRs in alignments with hundreds of diverse sequences from each protein, consistent with previous studies of intra-molecular contacts [3, 17], specifically when the alignments are deeper than they are long [19, 20]. We capture this rectangular quality in the statistic Neff/L, where Neff is the effective number of sequences as calculated by PSICOV [14] and L is the total number of columns in the pair of alignments. We observe similar trends when we use the number of sequences (N) or their phylogenetic diversity (PD), rather than Neff/L, to compare performance. The relationship between N, PD, and Neff is explored further in the Supplemental Text: *Diversity of sequences* and Supplemental Figures S10, S11 and S21. The diversity of residues within the individual alignment columns that make up each pair is another important factor to consider, and is explored in the Supplemental Text: *Performance by column entropy categories*.

Both power and precision improve with increasing Neff/L for nearly all coevolutionary methods (Figure 1), in the HisKA-RR data set. However, for alignments with Neff/L < 1.0, power at FPR<5% and precision at FPR<0.1% both remain relatively low (<50%). Additionally, the performance metrics f_max_ and phi_max_ show that there are no score thresholds (i.e. the strictness of predictions) that achieve both high precision and power in alignments with Neff/L < ~3.0 (Supplemental Figure S1). Despite the smaller range in Neff/L values, these performance trends are also observed across the 32 alignments in [24] (Supplemental Figures S3 and S6).

**Figure 1:**
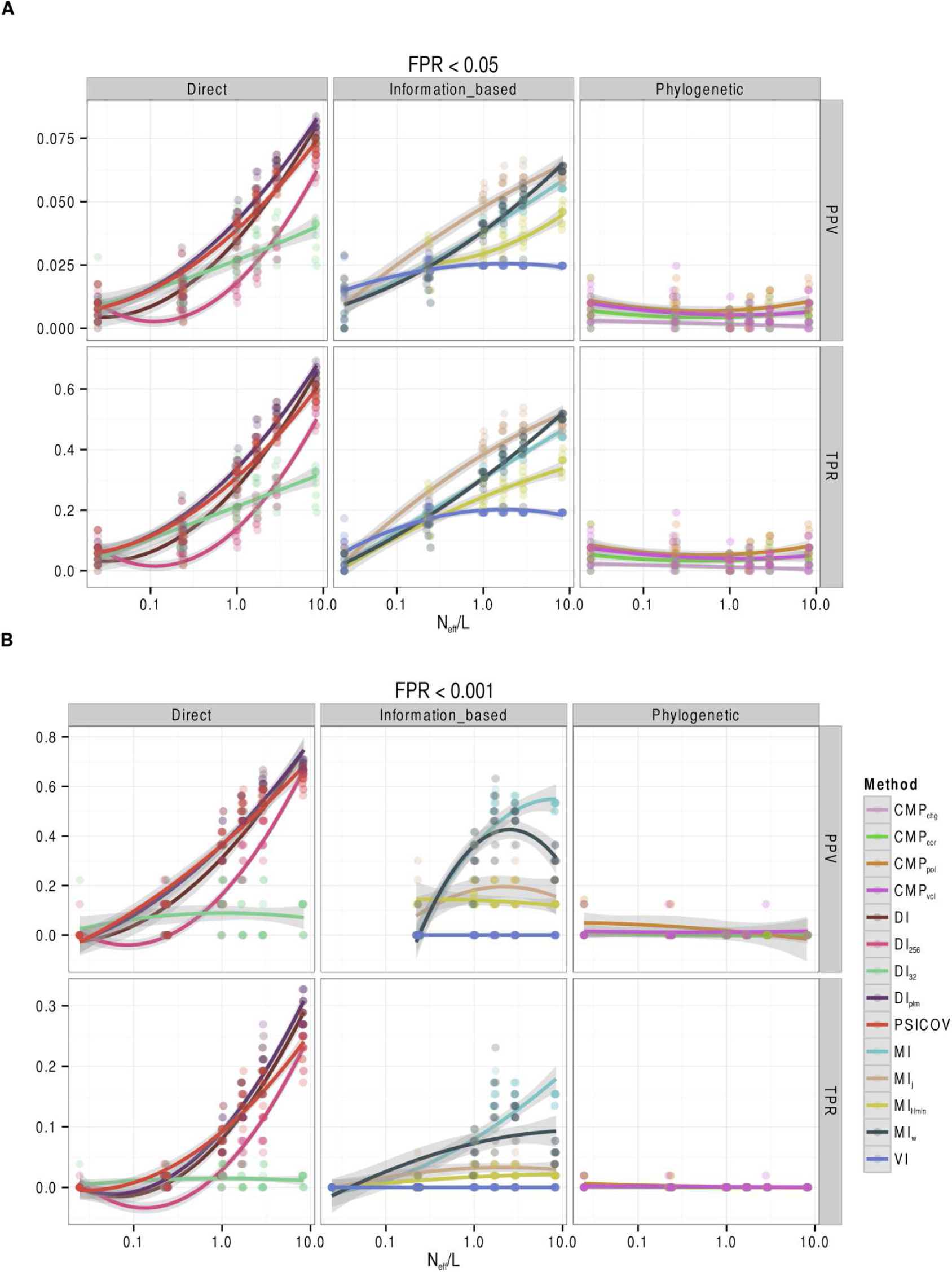
Coevolution statistics differ in their ability to detect residue contacts in HisKA-RR sub-alignments. Performance improves with larger, more diverse alignments. **A**: Power (TPR) and precision (PPV) at false positive rate (FPR) < 5%, **B**: at FPR < 0.1%. See Misc. Abbreviations and Table 1 for abbreviations.

In general, we confirm that coevolutionary methods that adjust for background phylogenetic signal through sequence re-weighting and/or average product correction (APC) (e.g., DI, DI_plm,_ and PSICOV) perform better than the phylogeny unaware mutual information (MI) based methods and the phylogeny aware approaches that explicitly use evolutionary models. In the HisKA-RR alignment, we observed two major exceptions to this trend when using the strictest definition for contacting pairs (i.e., requiring residue C*_β_* < 8Å coupled with biochemical evidence for specificity determination) (Supplemental Figure S2). First, the standard MI statistic is the most precise method for detecting contacting sites in alignments with Neff/L > 1.6 and FPR < 0.1%. Second, mutual information normalized by the joint entropy (MI_j_) has relatively high power in many scenarios and is the most powerful method for detecting contacting sites that are supported by experimental evidence at FPR < 5%. However, MI_j_ has drastically lower power at FPR < 0.1%. These findings suggest that MI is a reasonable choice if the goal of the analysis is to predict a small number of very high confidence contacts, whereas MI_j_ may be useful for detecting as many contacts as possible if a moderate FPR can be tolerated. These methods are both straightforward to compute, adding to their utility in these settings.

CoMap performance is an interesting case because, in contrast to DI, DI_plm,_ and PSICOV, it was not designed to find contacting residues. In the smallest alignments (5 sequences) we tested, it can have slightly better performance than the other methods. However, its poor performance in other alignments may indicate that it is identifying a set of coevolving residue pairs that partially overlap with contacting residues. It remains to explore whether CoMap can be used to prioritize residue pairs predicted by the other methods for functional assays.

Finally, we looked at the relationship between performance and the proportion of residue pairs that are contacts. Comparing across all 33 structures in our analyses, we observed the proportion of contacts is correlated with precision (Supplemental Figure S7). This means that most strongly coevolving residues in a protein pair are more likely to be physically interacting in co-crystal structures with larger interfaces.

### Choice of null distribution afects performance

The previous results show performance based on the known HisKA-RR structure. When applying the methods in our study in practice the structure usually is not known. One therefore uses a null distribution to control false predictions. Specifically, an upper quantile of the distribution of coevolutionary statistics in the absence of coevolutionary constraint is used as a threshold; one declares any pair of sites with a statistic exceeding the threshold a predicted contact. The goal is to minimize false predictions by predicting contacts only when statistics are much larger than expected by chance under the null distribution. A variety of null distributions are commonly used, including theoretical limiting distributions [5, 8], the observed empirical distribution (under the assumption that most pairs of sites are not coevolving) [39] and parametric, semi-parametric, and non-parametric bootstrap distributions [10, 40]. Theoretical and empirical nulls are more computationally efficient than bootstrap methods, which require simulating large data sets. The HisKA-RR interaction provides a framework for assessing the performance of these different approaches.

We used our sampled sub-alignments of HisKA-RR and the 32 alignments in [24] to compare the performance of two commonly used null distributions and to evaluate the sensitivity of each approach to alignment size. For each null distribution and coevolutionary statistic, we first employed the non-contact pairs of residues to assess if the FPR was truly controlled or not at given target FPRs of 5% and 0.1%.

The normal distribution is commonly used as theoretical null for mutual information and its normalized variants. Under this assumption, we standardized the coevolution scores to Z-scores and compared these to upper quantiles of the standard normal distribution (mean = 0, variance = 1). We then used the resulting upper-tail *P*-values (*P*_*normal*_) to predict contacting residue pairs. We found that nominal FPRs using this approach consistently exceed the target FPR across the range of Neff/L values in both the HisKA-RR sub-alignments and the alignments in [24] (Figures 2 and Supplemental Figure S4). In general, as Neff/L increases, the nominal FPR for Direct methods increases while it decreases in Information based methods. Nominal FPRs were up to twice to 20 times the target FPR for target FPRs 5% and 0.1% respectively. This suggests that either non-contacting residue pairs carry signals of coevolution (e.g., due to phylogeny, structural, or other evolutionary constraints) and/or that Z-scores of coevolution statistics have variance greater than one across non-contacting residues (e.g., due to an underestimated standard deviation across residue pairs resulting from within protein constraints or residues appearing in many pairs). Three of the four phylogeny aware CoMap methods controlled the nominal FPR below the target in all cases suggesting that the charge compensation analysis is predicting long-range residue interactions as well as contacts.

**Figure 2:**
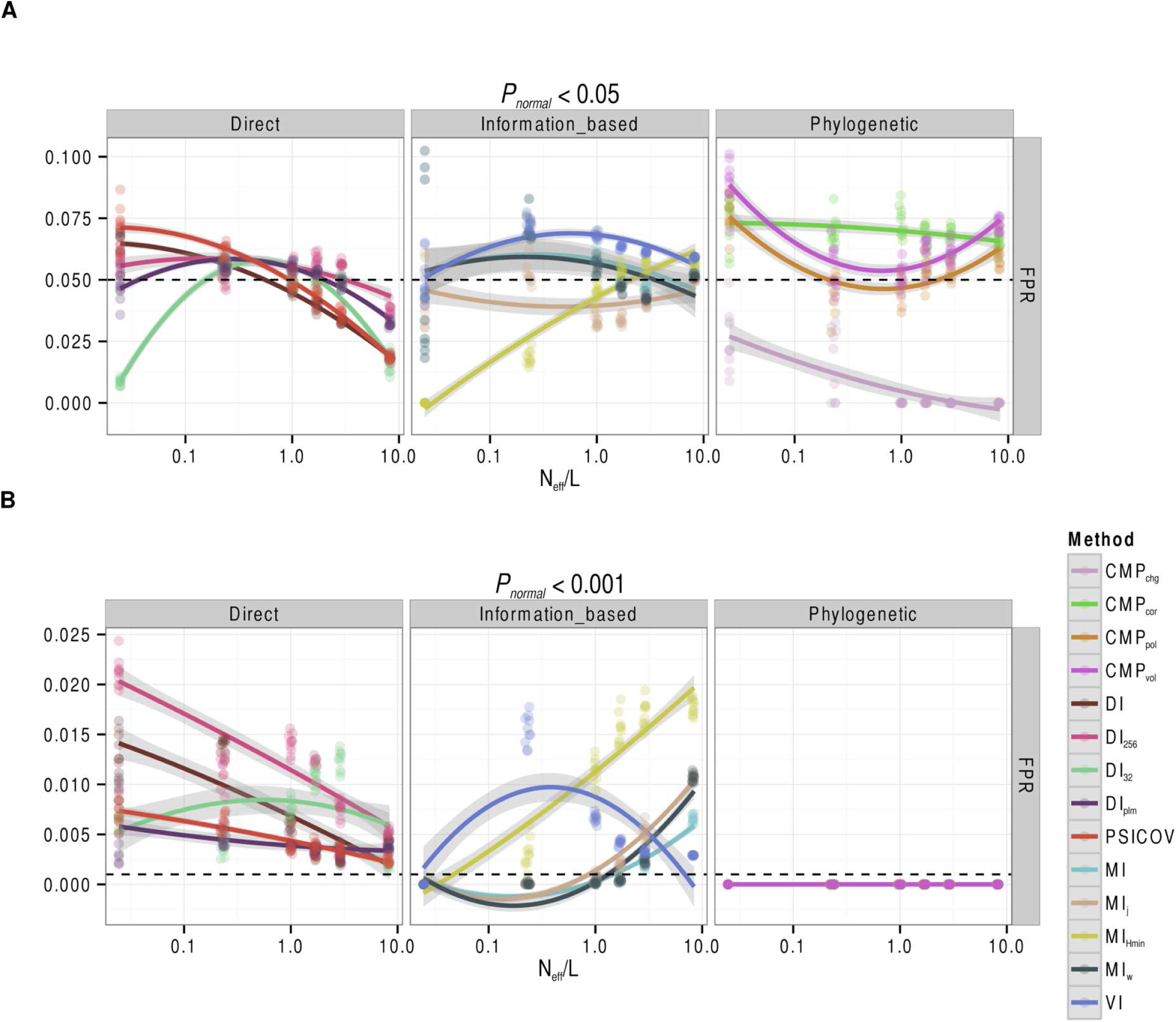
Commonly used null distributions for coevolution statistics’ null distributions often fail to control the false positive rate (FPR). **A**: Nominal FPRs for target FPR < 5%, **B**: target FPR < 0.1% (dashed lines) in the HisKA-RR alignments, assuming standardized scores have a standard normal null distribution, (i.e. using *P_normal_*). The phylogenetic methods control FPR at a threshold of 0.001, because they do not make any predictions at this significance level. See Misc. Abbreviations and Table 1 for abbreviations

Thus, while the normal distribution applied to standardized coevolution statistics can practically be used as a null distribution, we conclude that this approach results in elevated rates of false positive predictions, likely due to shared phylogeny or structural constraints affecting non-contacting residue pairs. A theoretical null (eg. noncentral gamma [41]) that is parameterized for individual column pairs may therefore be more appropriate.

Another choice of null distribution is the observed empirical distribution of the coevolution statistics. A *P*-value (*P*_*empirical*_) for a score *S* is simply the proportion of scores that are more extreme than *S*. This straightforward method can be easily applied with any statistic. However, it also assumes that no pairs of sites are coevolving and should therefore produce thresholds that are too strict when there are some coevolving sites in the data set (i.e., making it harder to predict real contacts). Contrary to this expectation, we found that the empirical null distribution—like the normal null distribution—produces nominal FPRs that exceed target FPRs (Figure 3 and Supplemental Figure S5). However, it is the Direct methods that best control the nominal FPR in both sets of alignments, marginally exceeding the target FPR in only a couple of cases. The Information-based methods fared well in the alignments in [24], however the HisKA-RR sub-alignments reveal that at Neff/L < 0.3, control of the FPR is lost, especially in MI_Hmin_. The Phylogenetic method that consistently exceeded the target FPR was the CoMap correlation analysis (CMP_cor_) which makes no assumptions regarding the biochemical properties of the amino acids. These results suggest that the empirical null distribution is not as conservative of an approach as one might expect from including contacting residue pairs in the null distribution. Although, it may suffer from some of the same effects that make the normal null distribution anti-conservative, such as shared phylogeny or structural constraints, alignments with very few sequences (eg. 5-50) have a limited number of possible scores which leads to ties in *P*-values between contacting and non-contacting residues.

**Figure 3:**
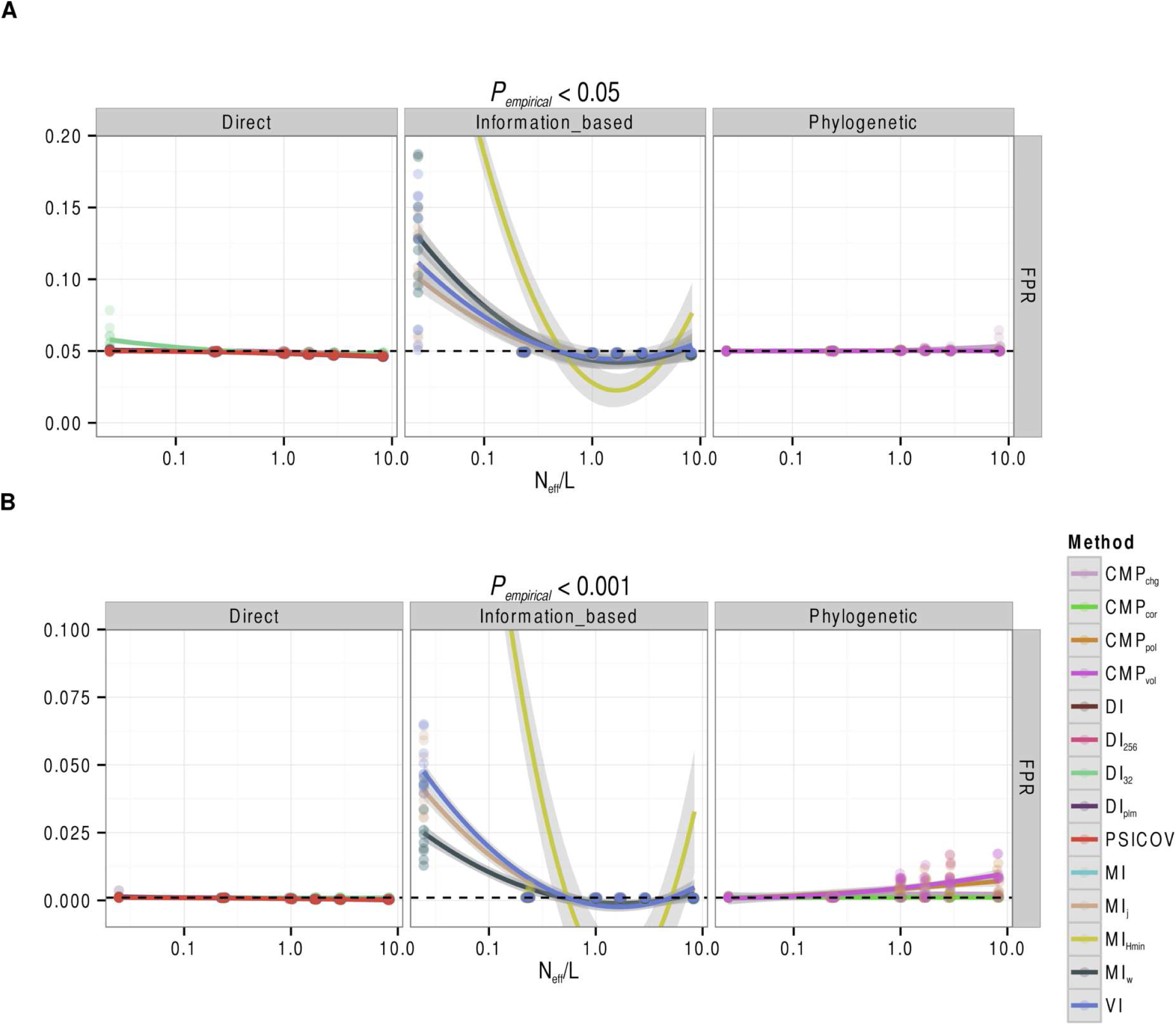
Commonly used null distributions for coevolution statistics’ null distributions often fail to control the false positive rate (FPR). **A**: Nominal FPRs for target FPR < 5%, **B**: target FPR < 0.1% (dashed lines) in the HisKA-RR alignments, using the empirical distribution of score ranks as the null distribution (i.e. using *P*_*empirical*_). See Misc. Abbreviations and Table 1 for abbreviations

These results are encouraging, but still leave us with the challenge of how to choose an appropriate *P*-value cutoff in a real analysis when the structure is unknown. Since our findings indicate that nominal FPRs exceed target FPRs with all three types of null distributions and nearly all methods, stricter *P*-value cutoffs than the target false positive rate seem warranted. But it is not clear how much stricter will be needed in any given alignment pair without additional information to guide such modifications (eg. incorporating alignment properties such as Neff/L into a model for each coevolution method). Hence, in most applications one must simply aim to control a target FPR, knowing that the true error rate is likely to be larger (Supplemental Figures S8 and S9). For this reason, the empirical null distribution may be the best choice to use as it controls error rates across the majority of alignment sizes, target FPRs, and coevolution methods (Figures 3 and S5) tested. As a rule of thumb, the empirical null overall controls the FPR for the Direct methods, however in small alignments (5 sequences or Neff/L < 0.3) it can be up to 1.5 times the target FPR.

### Cross-Species Case Study 1: Applying coevolution methods to Vif-A3G identiies some residues known to afect host-virus interactions

Viral infectivity factor (Vif) is a lentiviral accessory protein whose primary function is to target the antiviral cytidine deaminase APOBEC3G (A3G) of its mammalian hosts through ubiquitination. Because the two protein families are in an evolutionary arms race [42, 43], we hypothesized that they would be an informative example for exploring the utility of coevolution methods in host-virus protein pairs (i.e., inter-protein, inter-species interactions). This is a novel application of coevolution analysis, which has primarily been applied to residues within a protein or between pairs of proteins in the same genome.

A major challenge in performing coevolutionary analysis on cross-species protein pairs is acquiring appropriate data, including paired alignments and protein structures for validation. For Vif-A3G, we were able to identify 16 pairs of sequences (Neff = 10.0) from different primates (A3G orthologs) and their lentiviruses (Vif orthologs) in public databases (Table S2). Our benchmarking results on HisKA-RR indicate that such small protein families push the useful limits of the coevolution statistics we tested (Neff/L = 0.014). The low sequence diversity of A3G (Neff = 3.04) within primates compared to Vif (Neff = 11.3) within primate lentiviruses also presents challenges. Hence, we expect coevolutionary analysis to potentially have limited power in this scenario. To quantitatively evaluate performance, requires validated Vif-A3G interactions. The structure of Vif in complex with A3G has not been solved. However, biochemical assays have solidly identified regions important for binding and ubiquitination along the individual reference sequences of HIV1 Vif [44–47] and human A3G [48, 49] (Table S3). For this analysis, we therefore take the residues in biochemically-validated regions to be *positives* even though they might not be contacts (ie. C_β_ distance ≥ 8Å), and assume that all remaining residues are *negatives,* even though other sites (including sites deleted in these reference sequences) are possibly involved in the interaction. While further experimentation is needed to understand the relationship between functionally important sites and the structure of the protein interaction, as well as the effects of mutations in these sites on the fitness of lentiviruses, we explore whether any clues can be identified in the limited data that describes the coevolutionary history of the Vif-A3G residues.

First, we computed coevolutionary statistics for all Vif-A3G residue pairs and evaluated how well the statistics pinpoint the *positive* functionally important residues compared to *negatives*. For this evaluation, we used the empirical distribution of scores as a null distribution to determine statistical significance (i.e., *P*_*empirical*_) because they have lower false positive rates across Neff/L values at strict significance thresholds. Because the positives and negatives are single residues in each sequence instead of inter-protein residue pairs, we summarized *P*_*empirical*_ for each residue by assigning it the most significant *P*_*empirical*_ across all inter-protein pairs to which it belongs, and then explored the Vif and A3G results individually. From our benchmarking on the bacterial data sets, we know that significance thresholds that control the FPR vary by method and Neff/L, and that strict thresholds that yield very low (~2-3%) power are typically needed to control FPR in small alignments. We therefore chose to identify a significance threshold for each method that maximizes precision on the known functional sites in each protein. Then, we estimated power and FPR at these thresholds.

On Vif, with the exception of CMP_cor_ and DI_32_, the maximum precisions for each method ranged from 9 to 20% (i.e. only one or two residues out of ten predicted to be *positives* are truly *positives*)(Supplemental Figure S14). At these precision-optimized thresholds, MI_j_ and MI_minh_ predict almost every Vif residue to be coevolving; a stricter threshold would not result in a lower proportion of incorrect predictions. In contrast, the precisions for CMP_cor_, CMP_pol_, and DI_32_ are the highest (20%, 40%, 100% respectively). However, this comes at the cost of making the fewest number of predictions with the latter only making a single prediction. For these methods, less strict thresholds are needed to identify a greater proportion of *positives* at the cost of increasing the proportion of false discoveries. Across all methods, low f_max_ and phi_max_ values (0.26 and below) suggest there are no significance thresholds that balance power and precision for this data set.

We observed similarly low performance on A3G (Supplemental Figure S16). Encouragingly, we note that positions 128-130 are correctly identified by multiple methods (Supplemental Figure S12B). Residues at position 130 (e.g., D vs A) are highly likely to be adaptations that conferred species-specific resistance to Vif-induced degradation in Old World Monkeys 5-6MYA [42, 43]. Position 128, that also provides species-specific resistance, is thought to be more recent [42, 43, 50]. While these coevolution methods alone may not yet be accurate enough to identify functional residues, they potentially enhance other evolutionary analyses. For example, of the many Apobec sites under positive selection [43], it is reasonable that lentiviruses are more likely shaping the evolution of those sites that coevolve with Vif than sites that coevolve with other viral or virus-like agents.

Secondly, we visualized the localization of Vif residues predicted to be coevolving with A3G on a partial structure of Vif in complex with cofactors utilized for protein ubiquitination [51] (Figure 4). In [51], the authors are able to see that a critical subset of the Vif *positives* is solvent-exposed. We reevaluated performance with only these residues as the *positives* (Supplemental Figure S15). There is poor precision to identify the putative solvent-exposed interface among the methods; CMP_cor_ at 50% and CMP_vol_ at 10% are the only methods with precision >6%.

**Figure 4:**
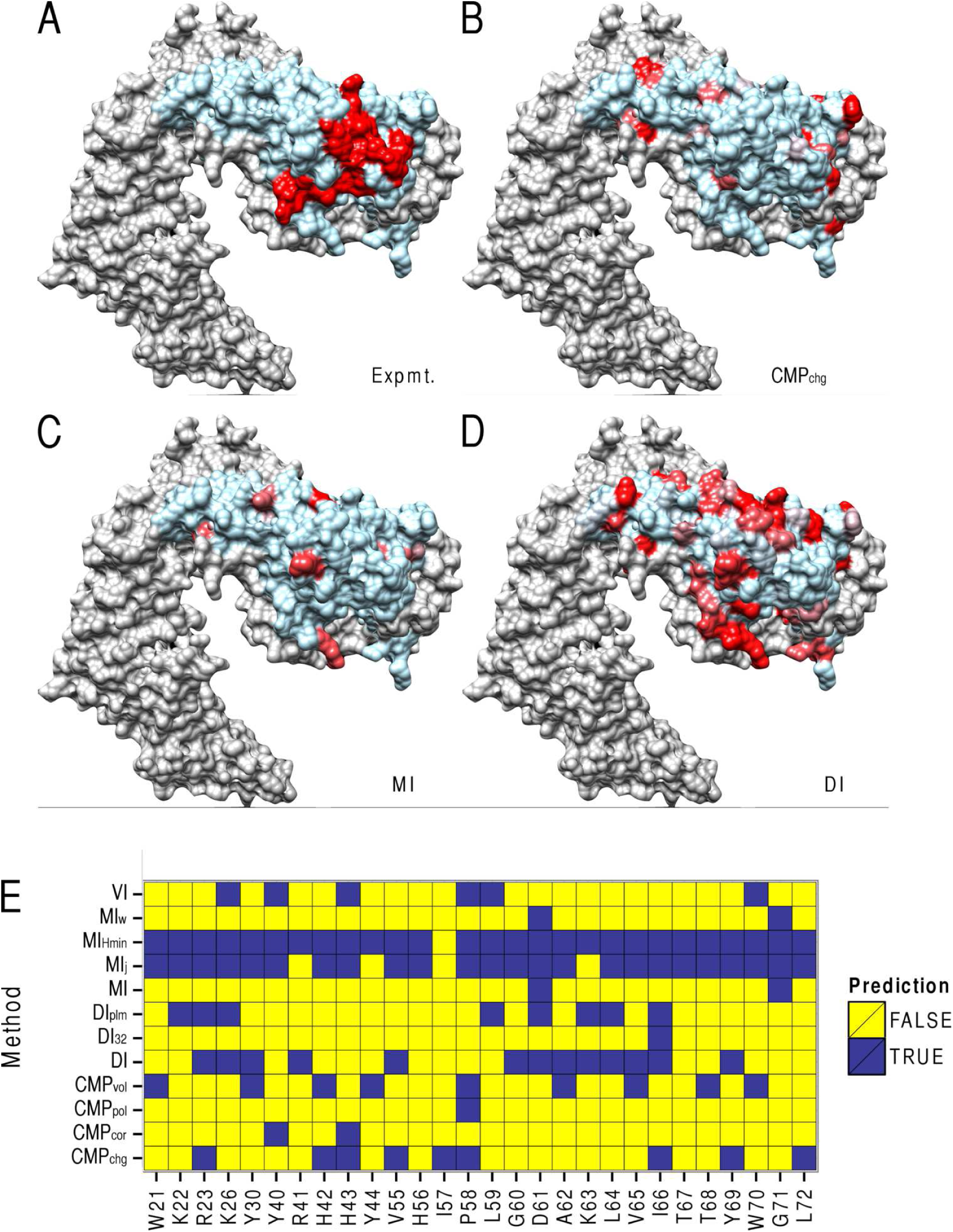
HIV1 Vif (light blue) in complex with co-factors (grey) sans APOBEC3G (A3G) (PDB ID: 4N9F). Residues in red are predicted to be coevolving with A3G optimizing precision (PPV) using **A**: previously known essential residues, **B-D**: predictions using CMP_chg_, MI, DI respectively. **E**: Few Vif residues previously known to interact with A3G are correctly predicted by more than four methods and none by methods in all classes of methods (Information-based, Direct, Phylogenetic). See Misc. Abbreviations and Table 1 for abbreviations.

Our analysis of the Vif-A3G interaction confirms that power to detect functionally important residues in each protein family is also low in inter-protein analyses between species, even though it is plausible that an arms race between lentivirus and mammal would give rise to stronger signals of coevolution compared to background. It is important to consider that perhaps the positions we considered *positives* may not all be of equal evolutionary importance across primates. Interfaces may be gained or lost and the rapid evolution of the two proteins likely produces many alternative solutions to maintaining an antagonistic interaction. There were many predicted positions that were not in the *positives* and further systematic validation and more comprehensive sequencing of lentiviruses and primates is needed to determine which pairs of residues are actually in close proximity or functionally required for other reasons. Additionally, there appears to be some level of complementarity in the predictions made by VI and MI_minh_ and the CMP methods, which measure different biochemical trade offs between coevolving residues. This strengthens the rationale for integrating methods to better predict interface residues experiencing potentially different evolutionary constraints (e.g., structural, catalytic activity, specificity). Coevolutionary analysis can help to generate and prioritize candidates for these experiments.

### Cross-Species Case Study 2: The interaction network of HIV and human proteins shows only weak evidence of coevolution across mammals

We sought to use inter-protein residue coevolution to refine a recently derived APMS protein-protein interaction network of the HIV-human interactome [33]. This study detected human proteins that interact with each HIV protein, either via direct physical contact or as members of complexes. Specifically, we hoped to use evidence of sequence coevolution to resolve direct versus indirect protein interactions amongst all human proteins measured to interact with each HIV protein. Secondly, we wanted to know if coevolutionary signals are strong enough to pinpoint key residues involved in the interfaces of any direct interactions.

For each protein in the HIV genome, we computed a multiple sequence alignment with all other sequenced immunodeficiency viruses that infect mammals with sequenced genomes. Similarly, we generated a multiple alignment of each human protein with the sequences of its orthologs from any mammal with a sequenced immunodeficiency virus. This produced pairs of host-virus protein alignments with up to six immunodeficiency viruses and their primate, feline, and bovidae hosts. For each pair of residues in a host-virus protein pair, we quantified coevolution using MIj and a semi-parametric bootstrap to calculate *P*-values (See Supplemental Text: *Simulating independently evolving pairs of alignments*). For each protein pair, we varied the significance threshold and computed the count of significantly coevolving residue-pairs. We then compared this statistic for interacting protein pairs from the APMS network versus a control set of 100 randomly chosen lentivirus-mammal protein pairs not included in the APMS network. We found that APMS detected interactions have only marginally more counts of significant signals of coevolution compared to non-interactions (best auROC = 0.541 at *P*_*bootstrap*_ < 0.0001), and therefore counts of coevolving residues are not sensitive enough to distinguish direct interactions or the residues involved in them for this set of virus and host proteins. Based on our benchmarking, we conclude that this lack of signal may result from low power due to the lack of sequenced lentivirus-mammal proteome pairs.

## Discussion

In this work we aimed to paint a picture of the performance of emerging methods to identify inter-protein contacts using coevolution and to identify properties of alignments where performance is expected to be best. As previously noted in intra-protein predictions [3, 9, 14], re-weighting of the sequences to account for the underlying phylogeny is important for inter-protein predictions as well, however as the comparison between MI_w_ and MI shows, it is important to tune the parameters controlling the re-weighting in cases where there are fast evolving alignment columns in an overall conserved protein family. Fortunately, methods that search for direct correlations—using a global statistical model for the sequence alignments—seem to be able to correct for the improper weighting (compare MI_w_ to DI). These methods are more precise at strict false positive rates than their counterparts especially when the alignments have Neff/L < 1.0. However, it may be beneficial to use a faster, MI-based method if the use case allows for a relaxed FPR and is concerned with power versus precision.

We also investigated the use of three null models to control the false positive rate. Counter-intuitively, a null model that explicitly models evolution independently for each alignment fails to control the false positive rate. We believe that our simulated alignments are systematically scoring too low because they fail to capture the correct amount of variation in the observed alignments, resulting in artificially significant *P*-values, except for when the effects of having small alignment sizes results in overly conservative *P*-values. Using a standard normal or the empirical distribution of scores as null models also failed to control the false positive rate, likely due to the correlation structure imposed by the shared evolutionary history of the residues, the distribution of evolutionary rates of the residues, or because asymptotic assumptions do not hold at small sample sizes. Thus, choosing an appropriate *P*-value cutoff in a real analysis when the structure is unknown and alignment depth is shallow still remains a challenge. However, we show that in diverse enough alignments the empirical null successfully controls the false positive rate for Direct methods. As a future direction, we suggest exploring theoretical null distributions that can be parameterized for individual alignment column pairs such as [41] or further improving protein evolution simulators to generate distributions of scores where the evolutionary rates are more similar between the null and alternate hypothesis.

A related problem to the one discussed here is to search a large set of protein pairs (within or between species) to determine which ones are interacting. In this setting, coevolution method performance is potentially more important than when predicting contacting residues for known interactions, because the search space will contain so many negatives (i.e., non-interacting pairs). A permissive *P*-value cutoff will lead to a large number of false positives and that may misinform investigators, while being too strict will lead to false negatives that keep potentially important findings hidden. While models exist that identify cutoffs based on benchmark data sets (e.g., Supplemental Figures S8 and S9, [24]), it would be interesting to understand why the parameters in these studies are appropriate and if they generalize to all protein-protein interactions. Ideally, we would like to understand what a null model teaches us about phylogeny-induced coevolution in the absence of structural inter- or intra-protein constraints. Another challenge for predicting interacting protein pairs from coevolutionary tests is how to summarize statistics for individual pairs of residues to produce a single score for a pair of proteins. Based on some preliminary investigations of these questions, we conclude that it is unlikely that cross-species interacting protein pairs can be accurately distinguished from non-interacting pairs on a genome-wide scale.

The progress of high-throughput interaction mapping highlights the need for continued refinement of inter-protein coevolution detection methods. Given that improper re-weighting of sequences can negatively affect power and the false positive rate, perhaps expanding Direct methods to independently obtain sequence weights for each alignment or using an evolution-based probabilistic weight (such as in CoMap or using phylogenetic logistic regression) for unusual variation in each column is a logical next step forward. Another important contribution would be to develop a generalizable null model that can help differentiate contacts when there are very few sequences available for protein families. Furthermore, investigating the correlations among the coevolution statistics themselves in inter-protein data sets could potentially disentangle structural from non-structural coevolutionary forces as well as serving to construct an ensemble method. Comprehensively sequencing orthologous pairs of protein families is a straightforward way to test the usefulness of these future contributions while simultaneously enabling current methods to perform to their fullest.

## Conclusion

We benchmarked 13 coevolution methods on 33 protein interactions with associated sequence alignments of varying depths. We conclude that coevolutionary analyses of cross-species protein-protein interactions is largely hindered by a lack of phylogenetically deep protein alignments for many proteins, and furthur demonstrate this in two example use cases involving HIV-human protein interactions. Additionally, we report that commonly used null distributions generally fail to control false positives in coevolutionary analyses, though errors are best controlled by the empirical null in large alignments.

## Materials and Methods

### Multiple sequence alignments

A master alignment of 8998 concatenated HisKA and RR sequences from [36] was graciously provided by the authors. From this alignment, aligned sequences were sampled uniformly (each sequence had equal probability of being sampled) to create sub-alignments with 5, 50, 250, 500, 1000, and 5000 sequences. We sampled 10 sub-alignments of each alignment size (number of sequences in sub-alignment), resulting in 60 total alignment pairs.

The alignments in [24] were downloaded from complexes section of the Baker lab website (http://gremlin.bakerlab.org/complexes/PDB/_benchmark/_alignments.zip) on Aug 29, 2014. The corresponding structures were downloaded from PDB and processed to obtain contacts between representative protein chains.

The CoMap implementation requires a preprocessing step to remove sequence redundancy (a data munging alternative to sequence weighting). This additional step was also necessary to prevent buffer underflow errors when evaluating likelihoods in very large input trees. Therefore, all alignments with more than 200 sequences were culled to contain the 200 most diverse sequences before being passed to CoMap. The sub-alignment used corresponds to the 200-leaf sub-tree that maximizes PD for each original input alignment and tree.

### Measuring coevolution

The coevolution methods used are listed in Table 1 and Table S1. Wrappers for the Direct methods are provided in coevo_tools to facilitate running from the command line. For methods in the plmDCA, mfDCA and hpDCA packages, MATLAB, or the MATLAB runtime executable is required as well as various MATLAB Toolbox dependencies and licenses.

### Evaluating coevolution performance

For each method, coevolution scores for pairs of amino acid positions were used to predict inter-domain pairs of amino acid residues that are close to each other in the representative co-crystal structure (PDB ID: 3DGE).

We define *positives* as pairs of alignment positions mapping to amino acid residues whose beta carbons (C_*β*_) are less than 8 angstroms apart in 3DGE. All other pairs of alignment positions are considered *negatives*.

We considered the following two alternative definitions of *positives*:

- Closest non-hydrogen atom-atom distance between residues is less than 6 angstroms
- C_*β*_ distance is less than 8 angstroms *and* at least one residue is mentioned as important in determining specificity of the HisKA-RR interaction in [52–56].

Residue pairs are predicted as coevolving if their scores or *P*-values are above a given threshold (eg. top 1%, *P* < 0.05) (Table S4).

### Phylogenetic diversity

Phylogenetic diversity (PD) is calculated as the sum of the branch lengths in a tree built from the concatenated multiple sequence alignment of both proteins. Trees were built using FastTree (version 2.1.7 SSE3) with options `-gamma -nosupport -wag`.

## Acknowledgements

We thank Martin Weigt for providing HisKA and RR alignments and for providing links to DCA source code. We also thank Julien Dutheil for help running CoMap correctly. This work was supported by a National Institutes of Health bioinformatics training grant, a UCSF Graduate Research Mentorship Fellowship, institutional funding from Gladstone Institutes, and a gift from the San Simeon Fund.

## Abbreviations

CoMap is abbreviated CMP in the main text and figures and CoMapP in supplemental figures. Effective number of sequences per column is abbreviated Neff/L. Phylogenetic distance is abbreviated PD. MI_Hmin_ appears as MIminh in figure legends. Precision (PPV) optimized metrics: ppvcut, ppvmax, ppvTPR, ppvFPR are the *P*_*empirical*_ threshold that maximizes PPV, said maximum PPV, power (TPR), and false positive rate (FPR) at said threshold.

## Authors’ contributions

AA carried out the analysis. AA and KSP designed the analysis and wrote the manuscript. All authors read and approved the final manuscript.

## Authors’ information

AA is a Bioinformatics graduate student at the University of California San Francisco in the laboratory of Dr. Katherine S. Pollard.

KSP is a Senior Investigator at the Gladstone Institutes and Professor of Epidemiology and Biostatistics at the University of California San Francisco.

## Supplemental Text

### A toolkit for inter-molecular coevolution analysis

In order to build a software suite for evaluating existing approaches to coevolution analysis, we first obtained implementations for a collection of intra-molecular coevolution software tools spanning the range of methods in the literature (Table 1). The coevolutionary methods in our analyses can be divided into two major groups, those that consider each pair of sites independently and those that consider pairs of sites in the context of a global statistical model for the multiple sequence alignment. Other methodological differences include the incorporation of two additional techniques that have been shown to improve performance, re-weighting sequences such that similar sequences contribute less to the final score [1] and applying an Average Product Correction (APC) to remove background noise and phylogenetic signal from “raw” coevolution statistics [2]. Of the methods we benchmarked, only CoMap (1) explicitly uses a phylogenetic model in its calculation of a coevolution statistic, (2) accounts for biochemical and physical properties of amino acid residues, and (3) reports a *P*-value based on internal simulation of independently evolving sites. In this benchmark we use the CoMap *P*-value as a statistic for comparison with other coevolution methods.

Our toolkit consists of three parts, (1) a collection of wrappers and post-processing utilities to facilitate running the coevolution programs from the command line and standardizing the diverse output formats into a single manageable file (https://github.com/aavilahe/coevo_tools), (2) an R package for calculating empirical and theoretical *P*-values and measuring performance (https://github.com/aavilahe/coevo_analysis_rpackage), and (3) scripts for visualizing coevolving residues on PDB protein structures (https://github.com/aavilahe/coevo_tools). We also implemented the canonical mutual information statistic, normalizations of mutual information in [3], and an information theoretic distance with desirable properties [4] not previously included in coevolution analyses (https://github.com/aavilahe/infcalc). Many of the coevolution methods we tested are computationally expensive, so we prepared our workflow to take advantage of multiprocessing workstations and high performance computing clusters.

We additionally designed our implementations to facilitate inter-molecular analysis by defining data structures, such as paired alignments and corresponding phylogenetic tree pairs, that accommodate analysis of multiple sequence alignment columns derived from two different proteins sequenced in potentially non-overlapping but matched sets of species. The matching of species is a key extension of standard gene family coevolution analysis to allow for interactome data analysis, where each sequence in one protein alignment is paired with one or more sequences in the second protein alignment (e.g., hosts and their viruses).

To extend CoMap-like *P*-values to other methods by simulating pairs of independently evolving protein alignments, we developed a semi-parametric simulation pipeline that combines software from the RAxML or FastTree, ANCESCON, Revolver, and HMMER3 packages [5–9] to estimate phylogenies from pairs of sequence alignments and then use these fitted models to generate large collections of protein sequence alignments that closely match observed protein families in alignment length, alignment size, phylogenetic diversity, amino acid composition, and domain architecture, in the absence of coevolution (see Methods). Our simulation pipeline is available at https://github.com/aavilahe/simulate_tools.

### Diversity of sequences

To investigate whether higher power in larger alignments results primarily from the number sequences per se or depends upon the diversity of the sequences, we compared the performance across alignments with different diversity values but the same number of sequences. We quantified diversity using phylogenetic diversity (PD) [10] and the effective number of sequences as calculated by PSICOV (Neff) [11] (Supplemental Figures S10, S11, S22, S23). For HisKA-RR sub-alignments, we found weak positive and negative relationships between the nominal false positive rate and PD for some methods in alignments with 5 sequences at given target false positive rates (Supplemental Figures S10, S11). While the range in diversity for such small alignments is small (PD: 7.5-11, Neff: 5), under the normal distribution, the false positive rate is better controlled in diverse alignments. However, under the empirical null, the Information-based methods do not control the FPR for these alignments and have larger false positive rates as diversity increases in these alignments.

One caveat of our HisKA-RR analysis is that (for computational reasons) we generated sub-alignments by random sampling and therefore only explored a range of phylogenies close to the typical diversity for each alignment size. The alignments in [12] provide a broader range of phylogenetic scenarios. Across these 32 protein pairs, we observe fairly strong correlations between Neff and performance (Figure S22, S23), although performance is quite variable at any Neff value. For example, the alignment pair with the highest Neff had the poorest performance while one with an intermediate value had the second best performance.

### Performance by column entropy categories

For a subset of methods, we measured the performance of the coevolution methods in pairs of columns with different rates of evolution. For each alignment size, the column entropies for each of the 10 HisKA and RR sub-alignments were aggregated and their median calculated. Then, for each sub-alignment, column pairs were binned into one of the following four categories:

1. above-median-HisKA-entropy + above-median-RR-entropy
2. above-median-HisKA-entropy + below-median-RR-entropy
3. below-median-HisKA-entropy + above-median-RR-entropy
4. below-median-HisKA-entropy + below-median-RR-entropy

Then for each category, the false positive rate, true positive rate, and precision were calculated, and the median performance is given in Supplemental Figures S17 and S18. Cutoffs and *P*-values that depend on the observed data are recalculated using only the column-pairs in each bin (eg. *P*_*normal*_, *P*_*empirical*_).

#### Most methods perform best on pairs of alignment columns with similar sequence variation in the two proteins

To explore the effect of substitution rate variation across sites in HisKA and RR, we parsed our performance results according to the entropy of the two alignment columns (one from each gene) in every pair of evaluated sites. For each alignment size, we split columns into below-versus above-median entropy separately for each gene, and then classified pairs of sites into the resulting four groups (see Methods). Then we computed power and precision separately for each rate category group. This analysis showed that faster evolving (i.e., above-median-HisKA paired with above-median-RR) contacts are generally the easiest to detect with coevolutionary methods. Dually conserved residues (i.e., low-HisKA paired with low-RR) (Supplemental Figures S17 and S18) are the next easiest to detect. We conclude that MI_w_’s drop in performance at 5000 sequences may be due to dually-variable columns being improperly reweighted. These analyses show that sequence variation quantitatively affects the accuracy of coevolution analyses, with most methods performing best when coevolving residue pairs have similar substitution rates.

### Simulating independently evolving pairs of alignments

In order to classify pairs of sites as coevolving or not coevolving using a semi-parametric bootstrapped null distribution, we calculated a *P*-value for the score at every pair of positions by comparing the observed score to the distribution of scores simulated for that pair under the null hypothesis (independent coevolution).

To simulate alignments, we used FastTree (version 2.1.7 SSE3) [6] to build maximum likelihood phylogenetic trees for the HisKA and RR protein families. We used hmmbuild from the HMMER3 package [9] (version 3.0 March 2010) to build a profile hidden Markov model (pHMM) for each family. We sampled amino acid residues from a first order Markov chain to generate an initial sequence for each family. Finally, we used Revolver (version 1.0) [8] to simulate 1000 alignments for each family independently. Revolver can simulate the evolution of a given root sequence that adheres to the domain constraints imposed by a pHMM, and preserves a similar phylogenetic history to the observed alignment. Revolver used the WAG substitution matrix and indel probabilities were set to zero in order to simulate constant length alignments. Gaps from the observed alignment were then overlaid on the simulated alignment. We automated this process in a pipeline available at https://github.com/aavilahe/simulate_tools.

A third type of null distribution is based on employing bootstrap methods to resample the observed alignment in ways that break coevolutionary correlation or to generate alignments from a model without coevolution. These approaches have the benefit that they directly account for phylogenetic effects in the null distribution and therefore have the potential to more accurately control FPRs but are computationally intensive and not suitable for all methods as they can greatly increase computational time. To explore this possibility, we implemented a semi-parametric bootstrap null distribution for the phylogeny unaware methods in the HisKA-RR sub-alignments as an example of this approach.

This null distribution aims to resemble the observed alignments in terms of substitution rates and patterns, but substitutions are generated independently in HisKA and RR and are therefore not correlated beyond any correlation induced by similarities in the phylogenies of the two gene families. Unfortunately, we found that *P*-values calculated using the bootstrap null distribution were heavily influenced by the error in simulating alignment columns with appropriate amino acid variation. Simulation error increased with alignment size, as did nominal FPRs. Residue pairs for which the bootstrap simulated alignment columns have too much sequence variation tend to have small *P*-values, regardless of whether or not they are contacting residues. Consequently, at a target FPR of 5%, the nominal FPR was not adequately controlled for alignments with more than 5 sequences (Neff/L = 0.02) for any method except PSICOV. Interestingly, PSICOV is the method least affected by the simulation error.

Recalculating the nominal FPR using only alignment column pairs that were moderately well simulated (no more than 250 of 1000 simulations were over or under conserved) showed much lower FPR for all methods except MI_Hmin_ (Supplemental Figure S19). MI and VI are controlled below a target FPR <5%. At a stricter target FPR < 0.1%, PSICOV, MI, and VI are the only methods with completely controlled FPR at all alignment sizes. MI_w_, DI, and MI_j_ are controlled in alignments with fewer than 1000, 500, 250 sequences respectively. Together these results suggest that the DI, MI_w_, MI_j_, VI are sensitive to the amount of variation in the simulated alignments, while PSICOV and MI_Hmin_ are more robust to predicting fast and slowly evolving columns. However, MI_Hmin_’s higher FPR suggests it is identifying coevolving residues that are not structurally close. Perhaps they may be part of an alternate network of evolutionarily important residues, for example “protein sectors [13]” that span more than one protein.

CoMap internally estimates *P*-values using a similar simulation approach. Nominal FPRs for CoMap methods, using their *P*-values directly, resemble those of the Information based approaches using the normal distribution as a null (twice to 20 times the target FPR). We conclude that it is very important for the evolutionary conservation of alignment columns in the bootstrap null distribution to closely match conservation levels in the observed data. Despite using currently accepted techniques for generating bootstrap distributions, we found that matching conservation levels this closely is challenging. This is an important problem for future research in the coevolution field.

1000 alignments were independently simulated for 8998 HisKA and RR sequences each.

First a phylogenetic tree for each alignment was built using FastTree (version 2.1.7 SSE3) with options `-gamma -nosupport -wag`.

The following steps were then automated in the simulate_tools pipeline:

1. Build profile HMM
2. Sample starting “root sequence” for simulation using first order Markov chain
3. Generate xml control file for Revolver

A. No tree scaling
B. Heterogeneous rates (alpha = 1, ncats = 9)
C. No indels
4. Run Revolver

*An example command for simulating 1000 RR alignments:*

~~~
runSimAli --tree RR.tree\
--outdir /path/to/output\
--num\_sims 1000 JobNameRR RR.phy
~~~

From these simulated master alignments, sequences corresponding to the observed sub-alignments were extracted to create a total of 60000 sub-alignments, each corresponding to one of the original 60 observed sub-alignments.

### Structure visualization

The Vif complex 4N9F was rendered using the UCSF Chimera package (version 1.81) from the Computer Graphics Laboratory, University of California, San Francisco (supported by NIH P41 RR01081) [14].

### Alternate theoretical null *P*_*gamma*_

[15] derive the noncentral gamma distribution for a mutual information estimator sufficiently accurate for when the true MI <0.2 bits. The shape and scaling parameters depend on the number of observations (eg. number of sequences in alignment) and number of realizations of the two categorical variables (eg. number of different residues with non-zero probability in each alignment column), and the noncentrality parameter is used to specify “true MI” under the null hypothesis.

**Figure S1:**
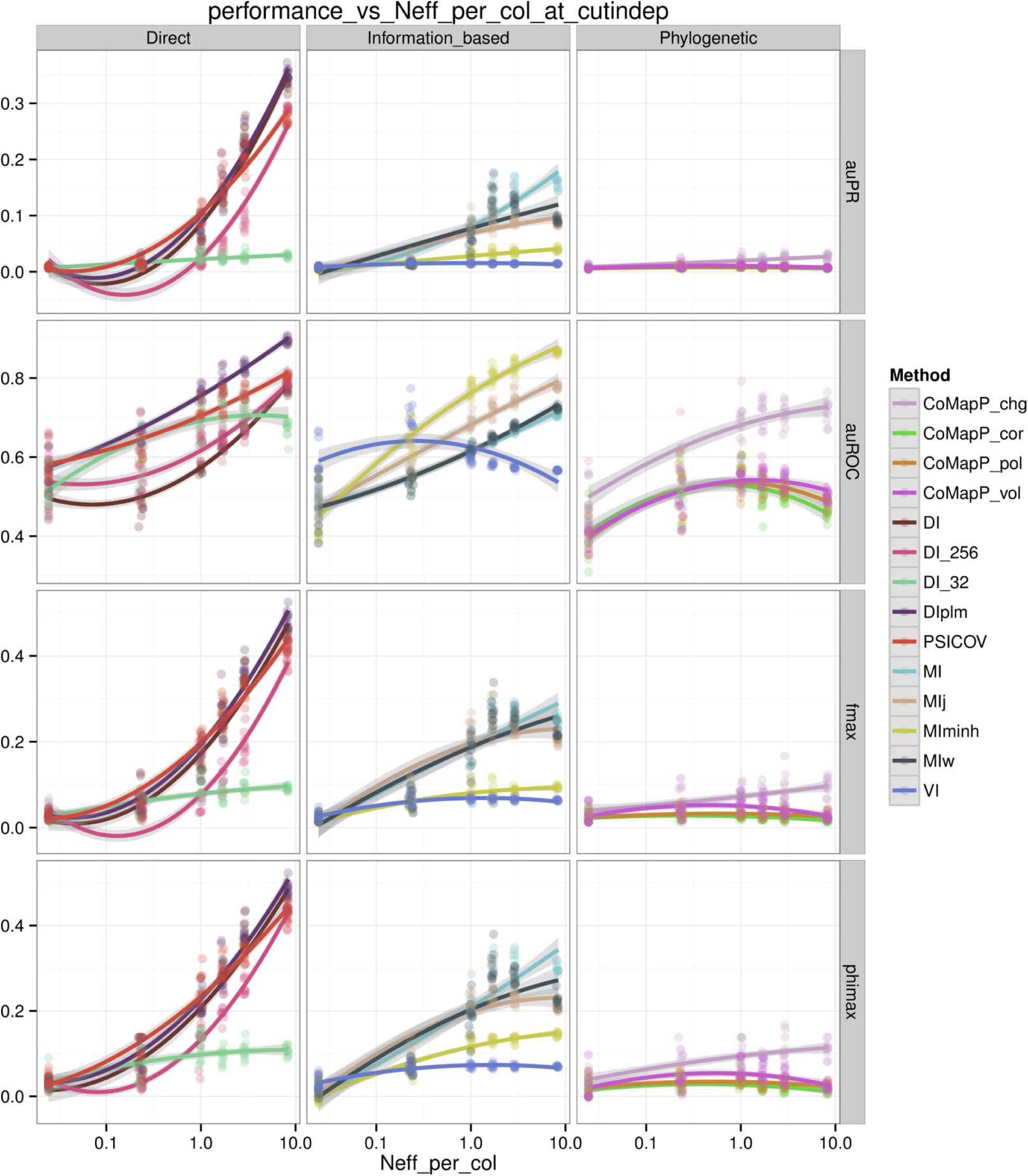
Threshold-independent performance metrics show that coevolution methods fail to achieve both high precision (PPV) and power (TPR) in HisKA-RR sub-alignments with Neff/L < ~3.0. See Misc. Abbreviations and Table 1 for abbreviations.

**Figures S2:**
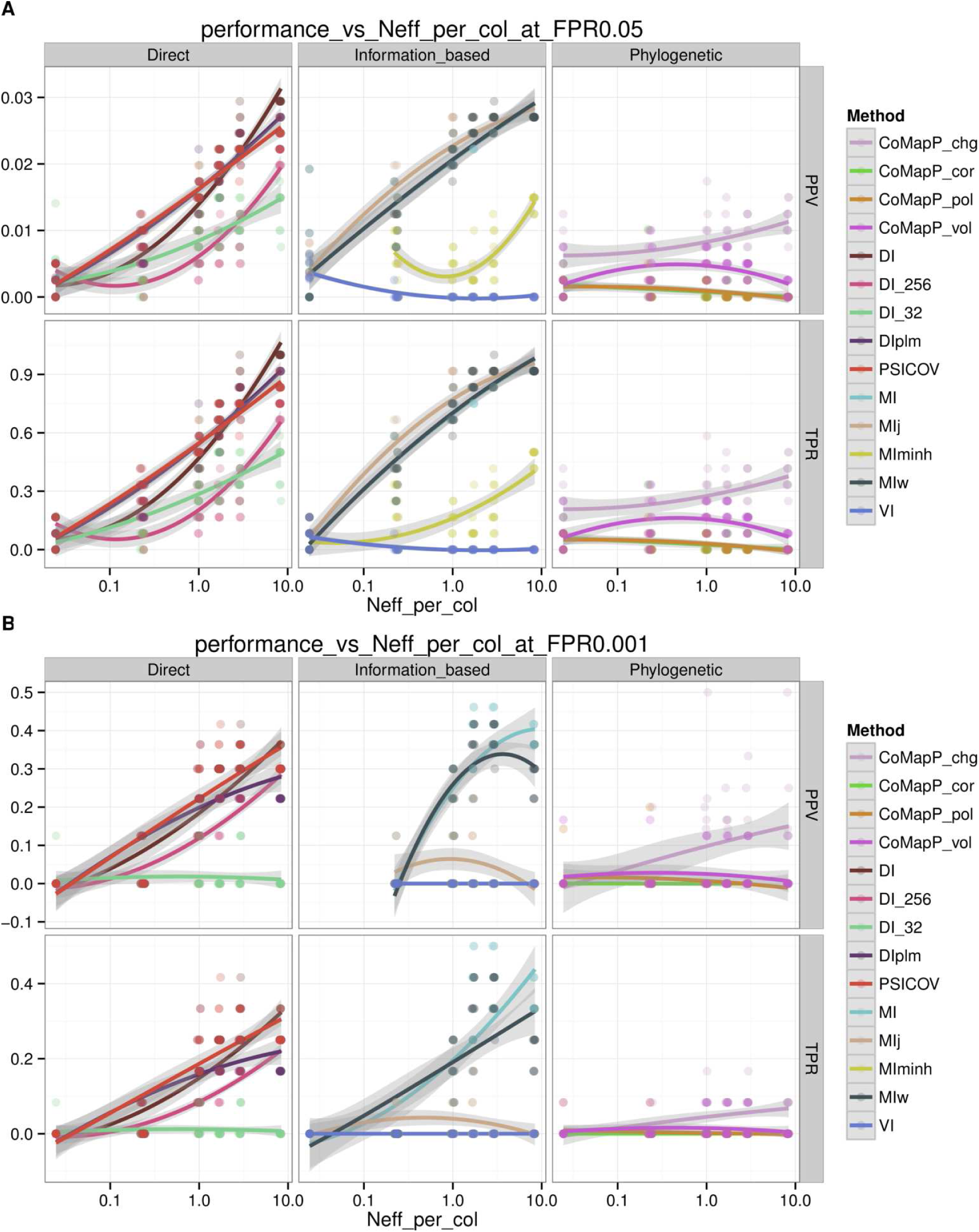
Power (TPR) and precision (PPV) at controlled false positive rates (**A**: FPR <5%, **B**: <0.1%) in HisKA-RR sub-alignments using a stricter definition for contacting residues that requires experimental evidence for specificity determination. See Misc. Abbreviations and Table 1 for abbreviations.

**Figure S3:**
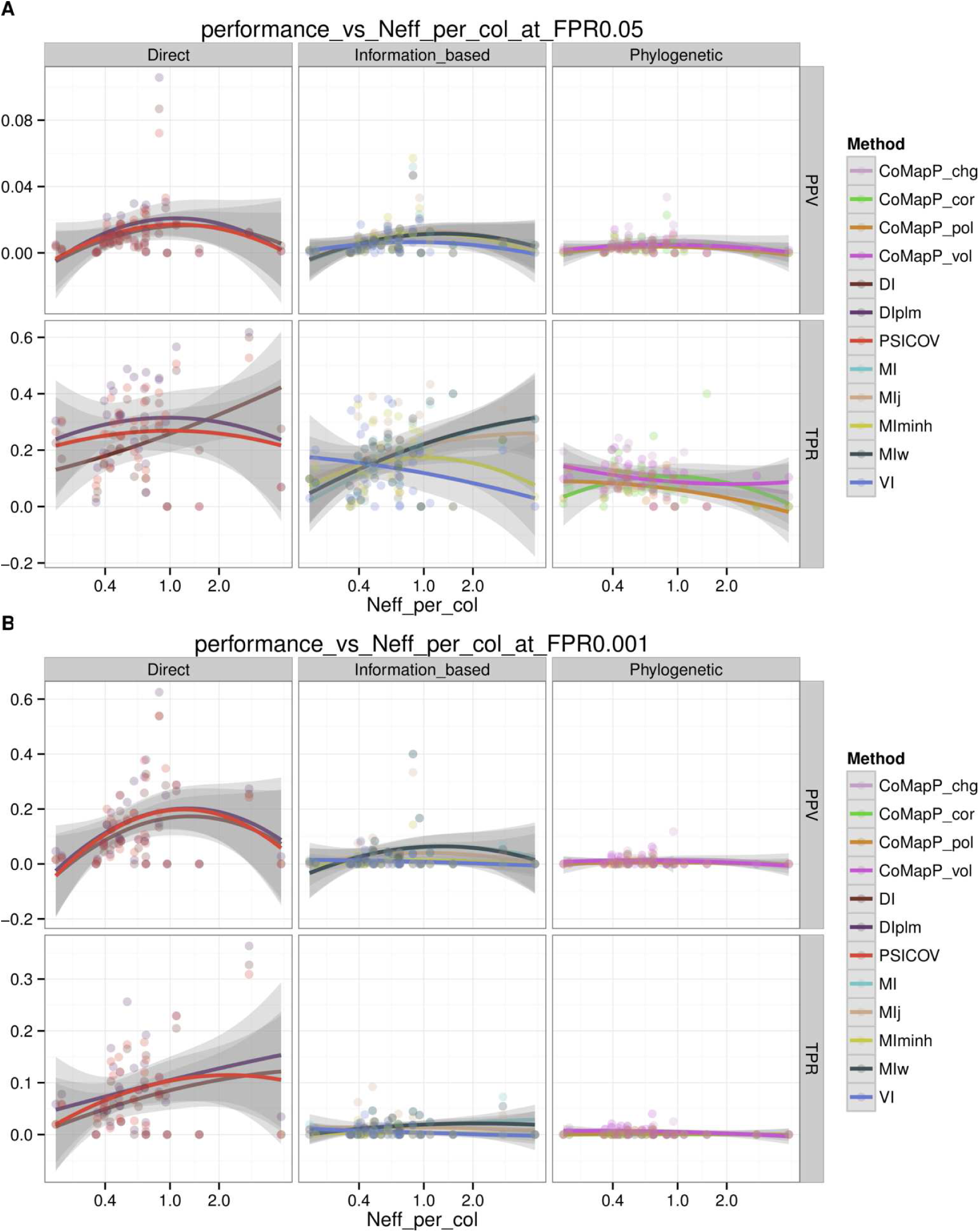
Coevolution statistics differ in their ability to detect residue contacts in the 32 alignments in [12]. Performance varies widely across the range of Neff/L values. **A** Power (TPR) and precision (PPV) at false positive rate (FPR) < 5%, **B**: at FPR < 0.1%. See Misc. Abbreviations and Table 1 for abbreviations.

**Figure S4:**
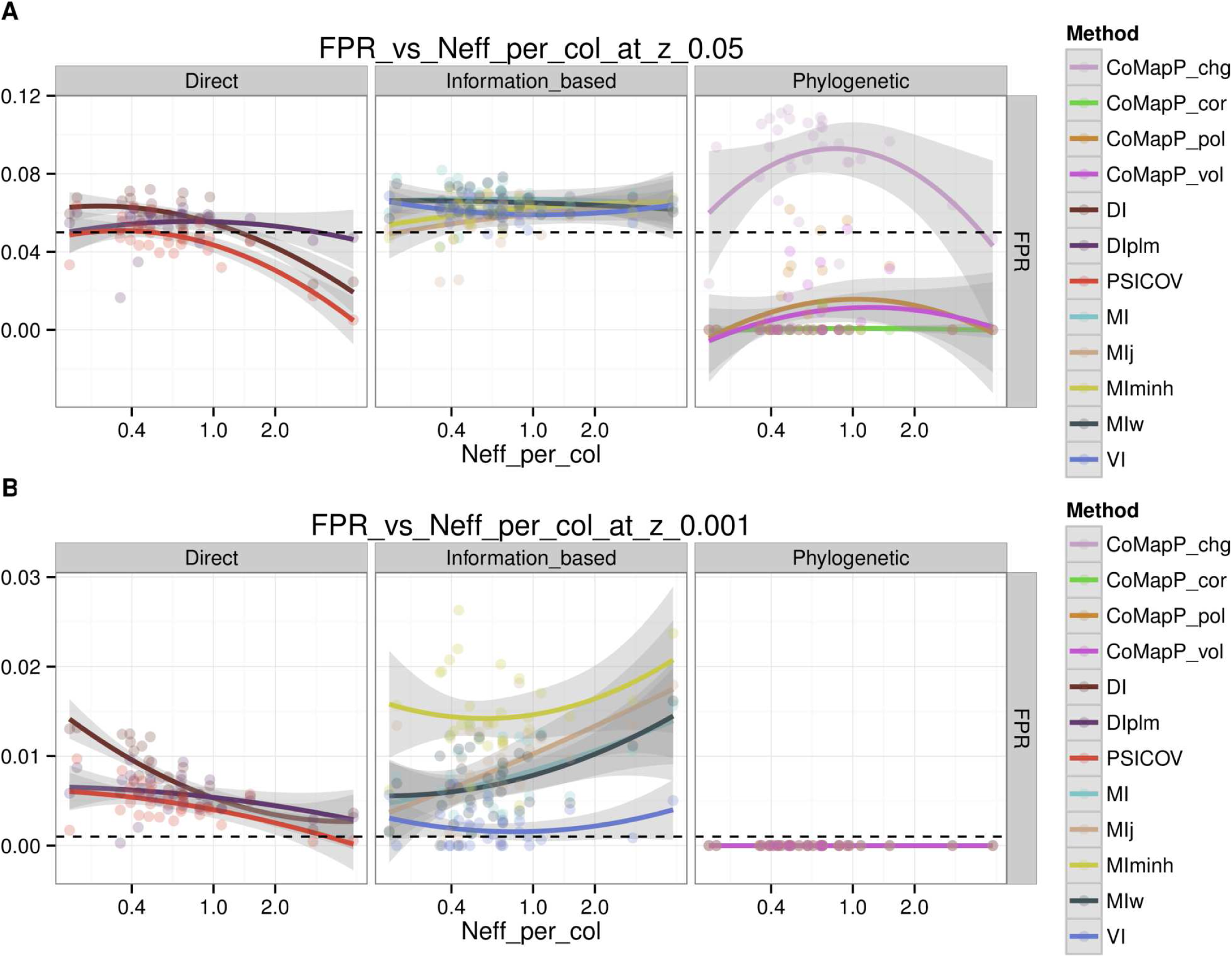
Commonly used null distributions for coevolution statistics’ null distributions often fail to control the false positive rate (FPR). **A**: Nominal FPRs for target FPR < 5%, **B**: target FPR < 0.1% (dashed lines) in the 32 alignments in [12], assuming standardized scores have a standard normal null distribution, (i.e. using *P*_*normal*_). See Misc. Abbreviations and Table 1 for abbreviations.

**Figure S5:**
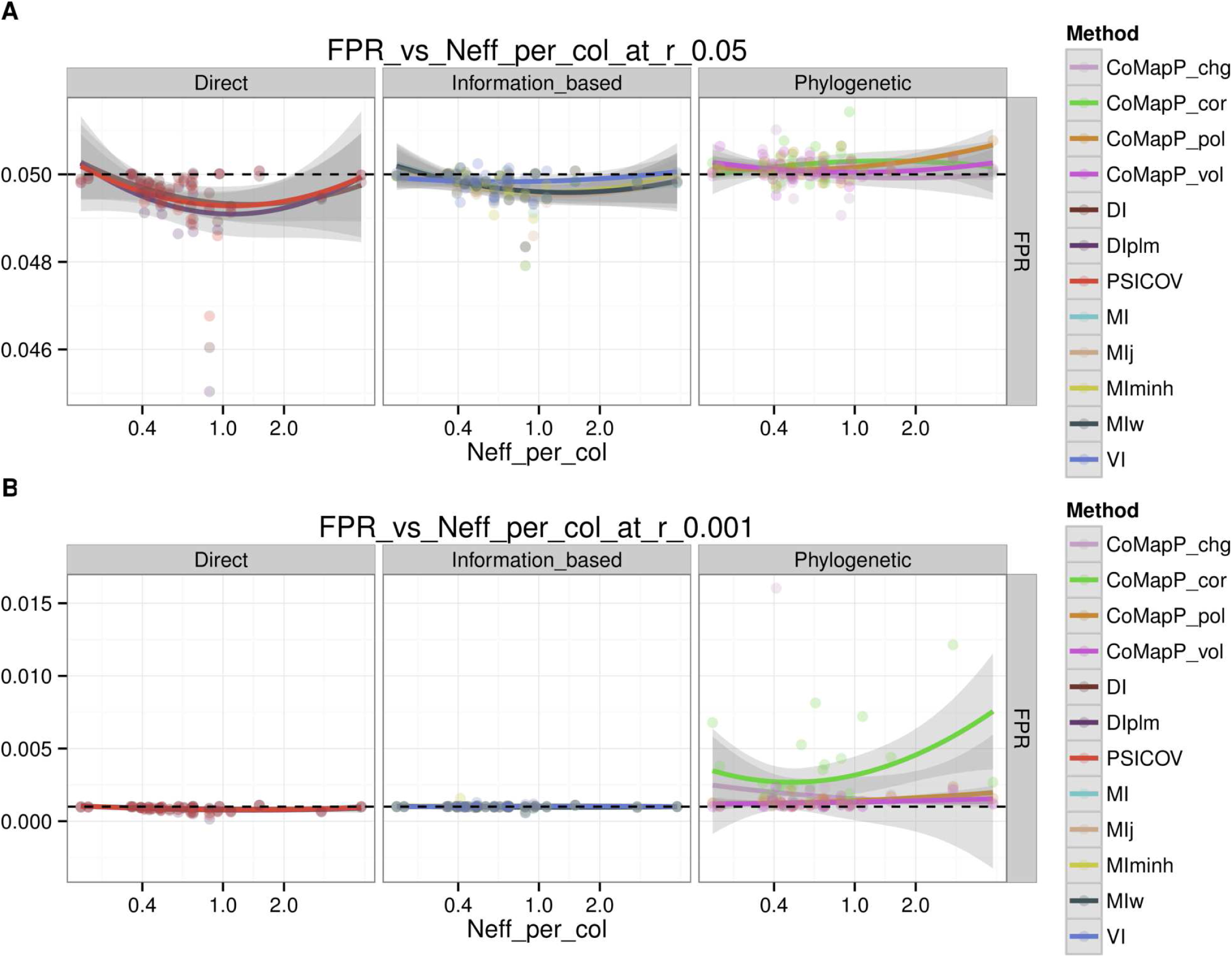
Commonly used null distributions for coevolution statistics’ null distributions often fail to control the false positive rate (FPR). **A**: Nominal FPRs for target FPR < 5%, **B**: target FPR < 0.1% (dashed lines) in the 32 alignments in [12] using the empirical distribution of score ranks as the null distribution (i.e. using *P*_*empirical*_). See Misc. Abbreviations and Table 1 for abbreviations.

**Figure S6:**
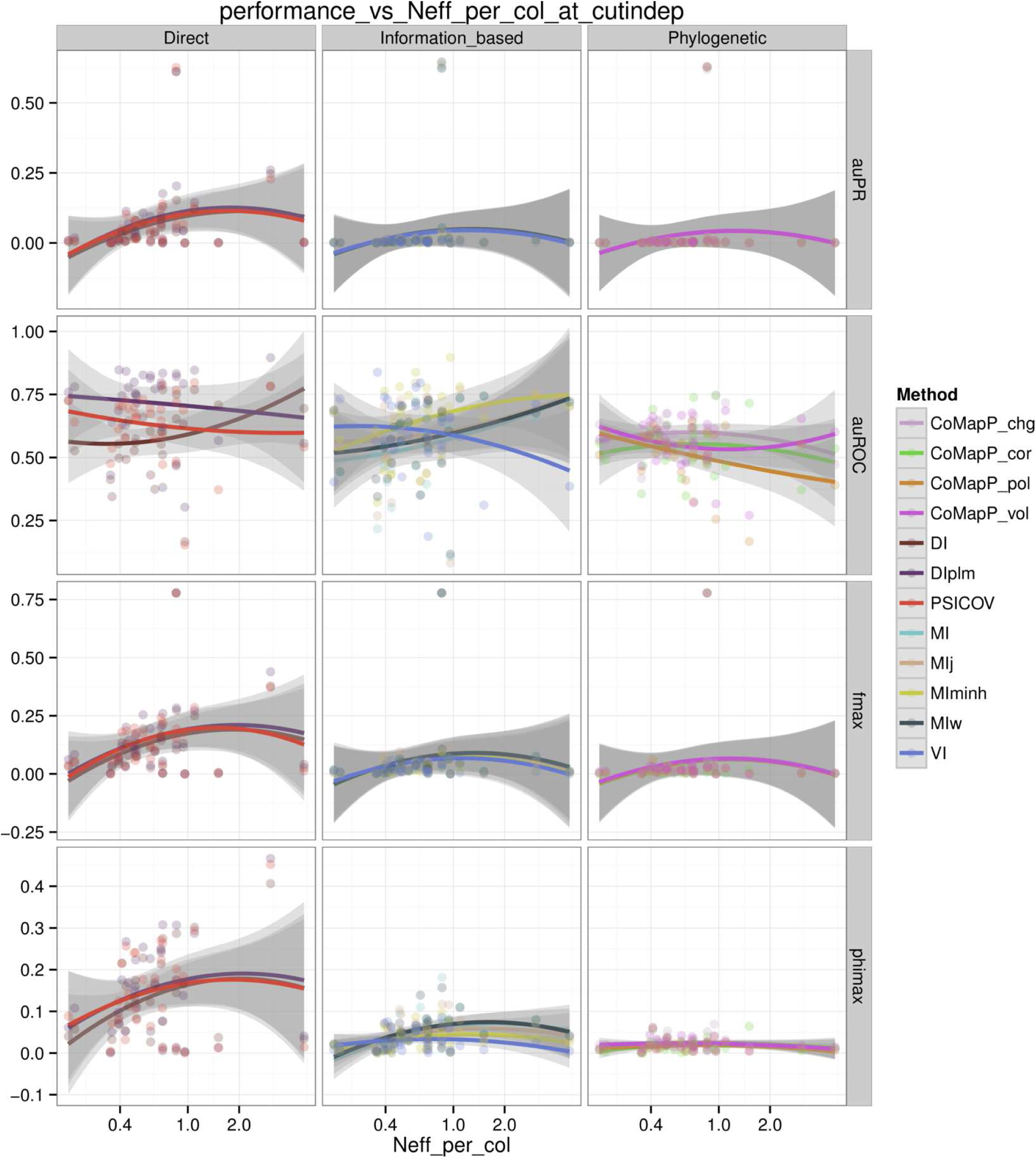
Threshold-independent performance metrics vary in the 32 alignments in [12] but trend upwards with increasing Neff/L. See Misc. Abbreviations and Table 1 for abbreviations.

**Figure S7:**
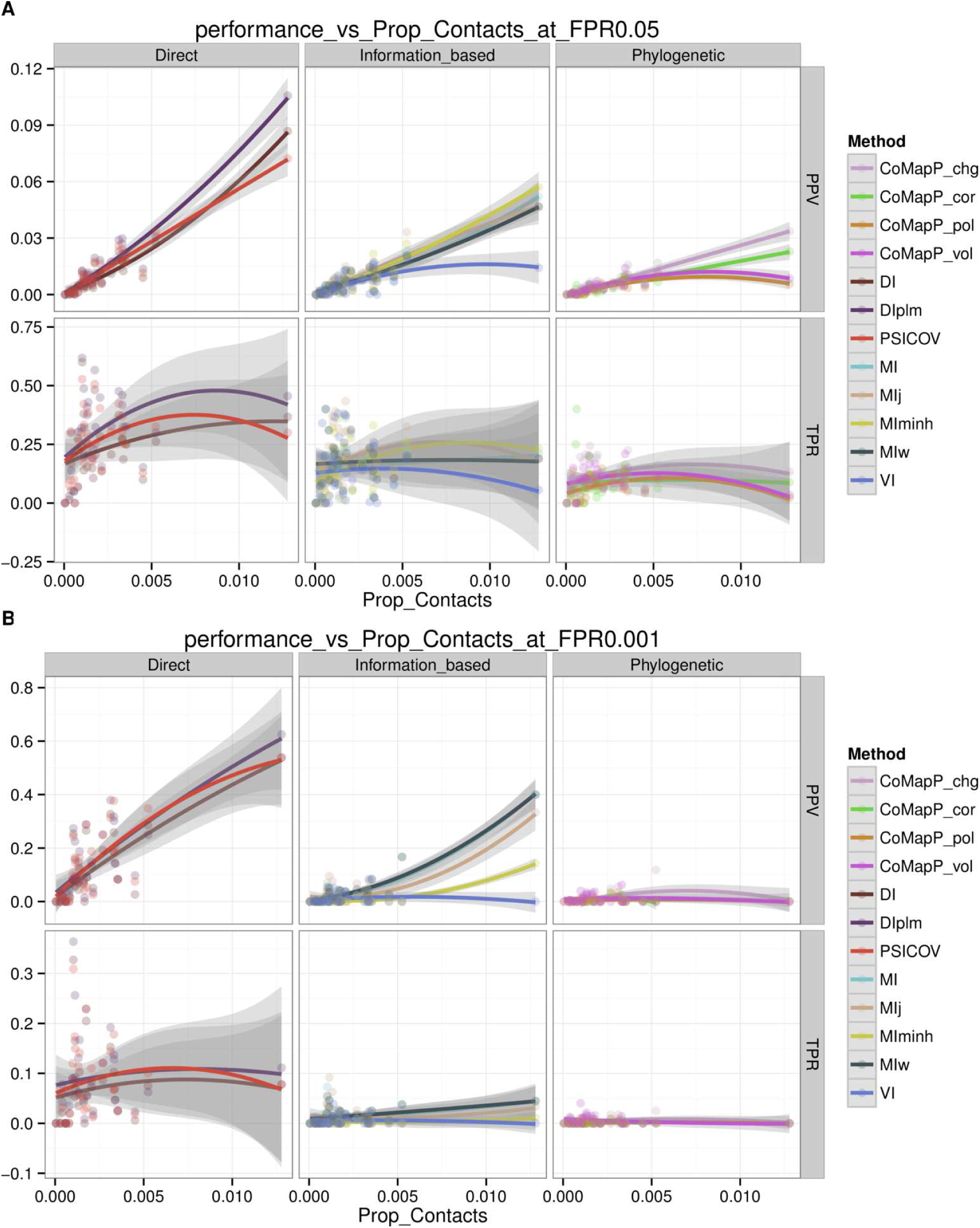
Precision (PPV) but not power (TPR) is positively correlated with the proportion of contacts in the 32 alignments in [12] at controlled false positive rates **A:** FPR < 0.1% **B:** FPR < 5%. See Misc. Abbreviations and Table 1 for abbreviations.

**Figure S8:**
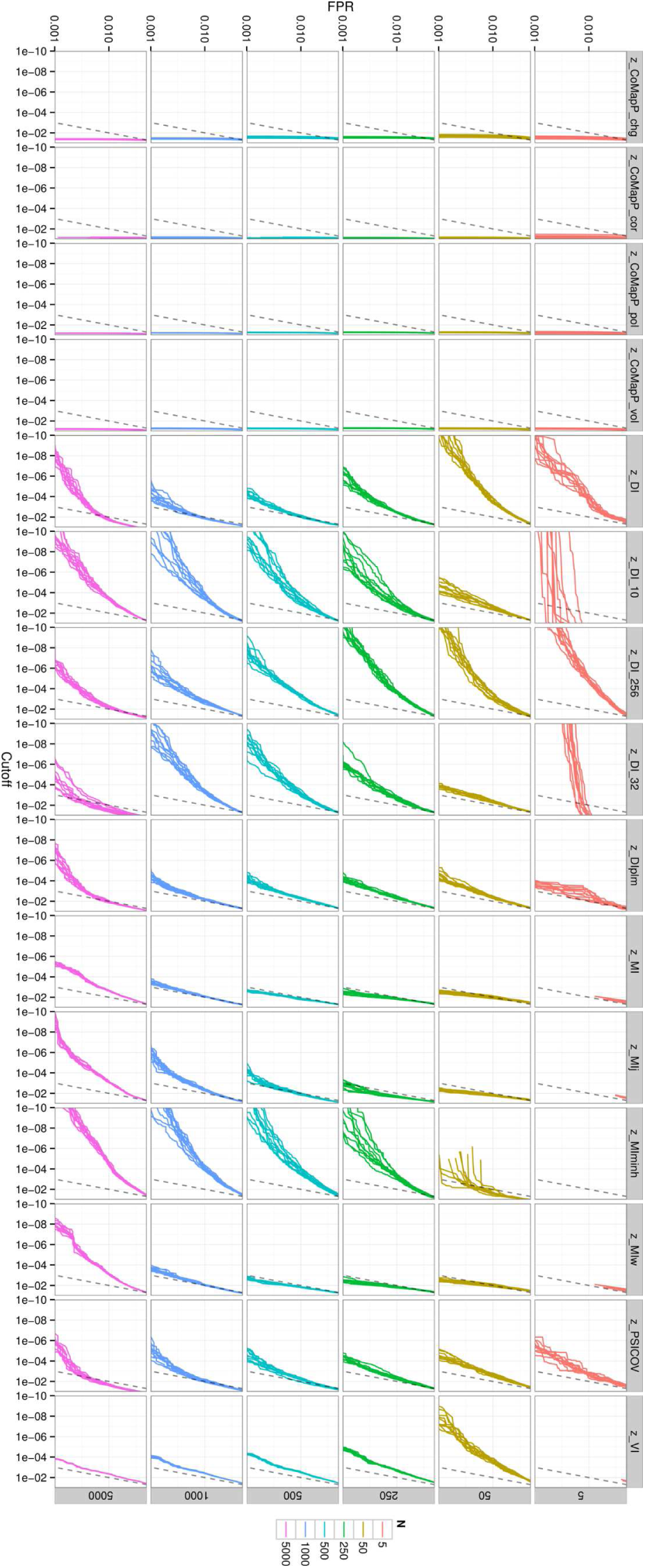
Nominal FPR at a given target FPR assuming a normal null distribution (*P*_*normal*_) for all 60 HisKA-RR sub-alignments. See Misc. Abbreviations and Table 1 for abbreviations.

**Figure S9:**
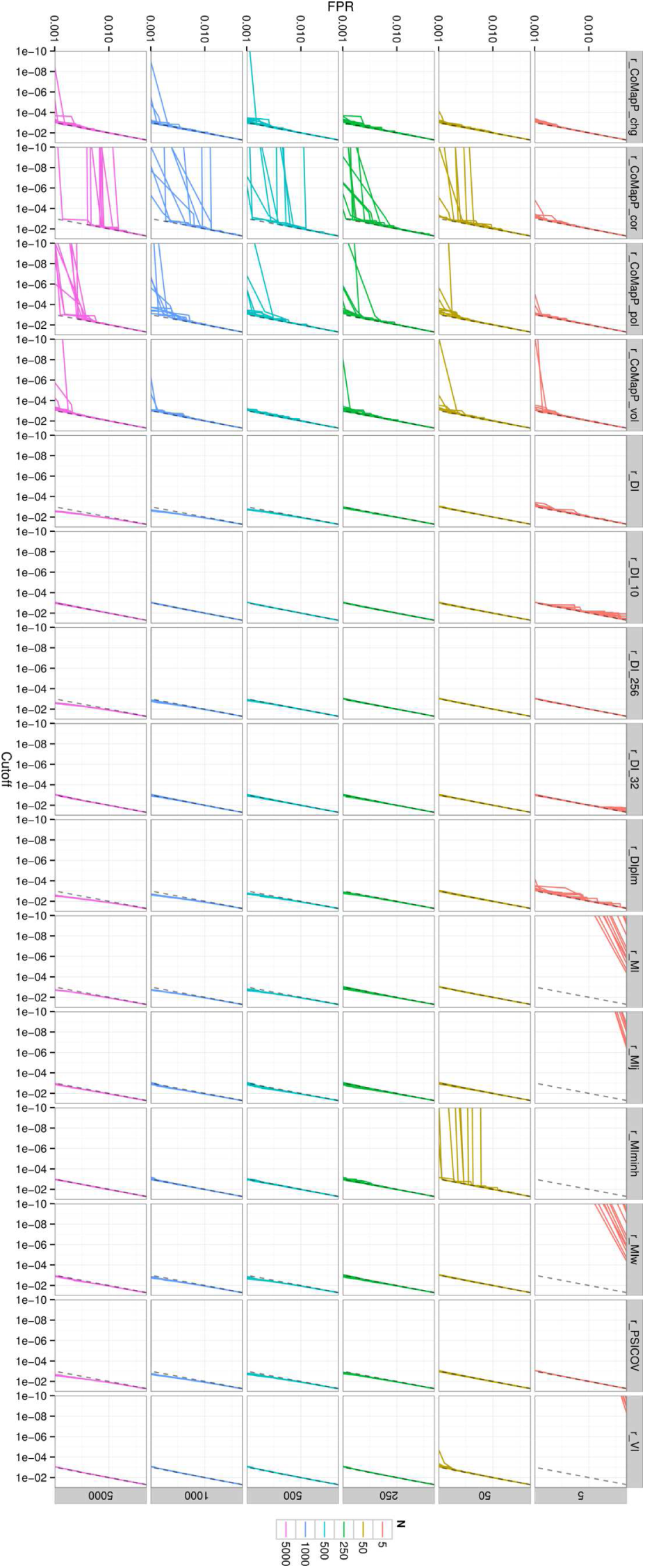
Nominal FPR at given target FPR assuming an empirical null distribution (*P*_*empirical*_) for all 60 HisKA-RR sub-alignments. See Misc. Abbreviations and Table 1 for abbreviations.

**Figure S10:**
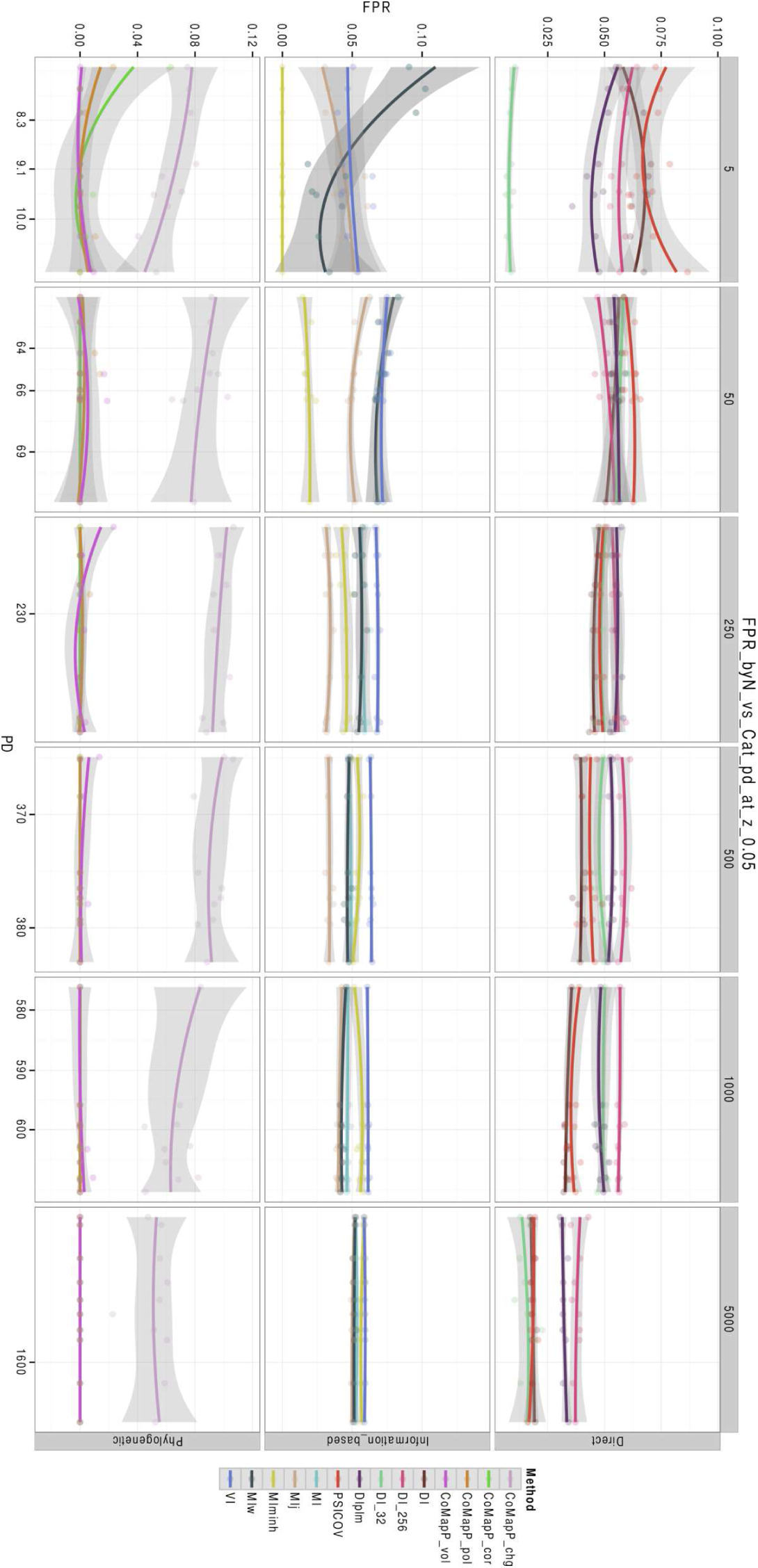
Sequence diversity may be important for controlling the false positive rate (FPR) in small alignments. Nominal FPR vs phylogenetic diversity (PD) at *P*_*normal*_ < 0.05. PD is the sum of branch lengths. See Misc. Abbreviations and Table 1 for abbreviations.

**Figure S11:**
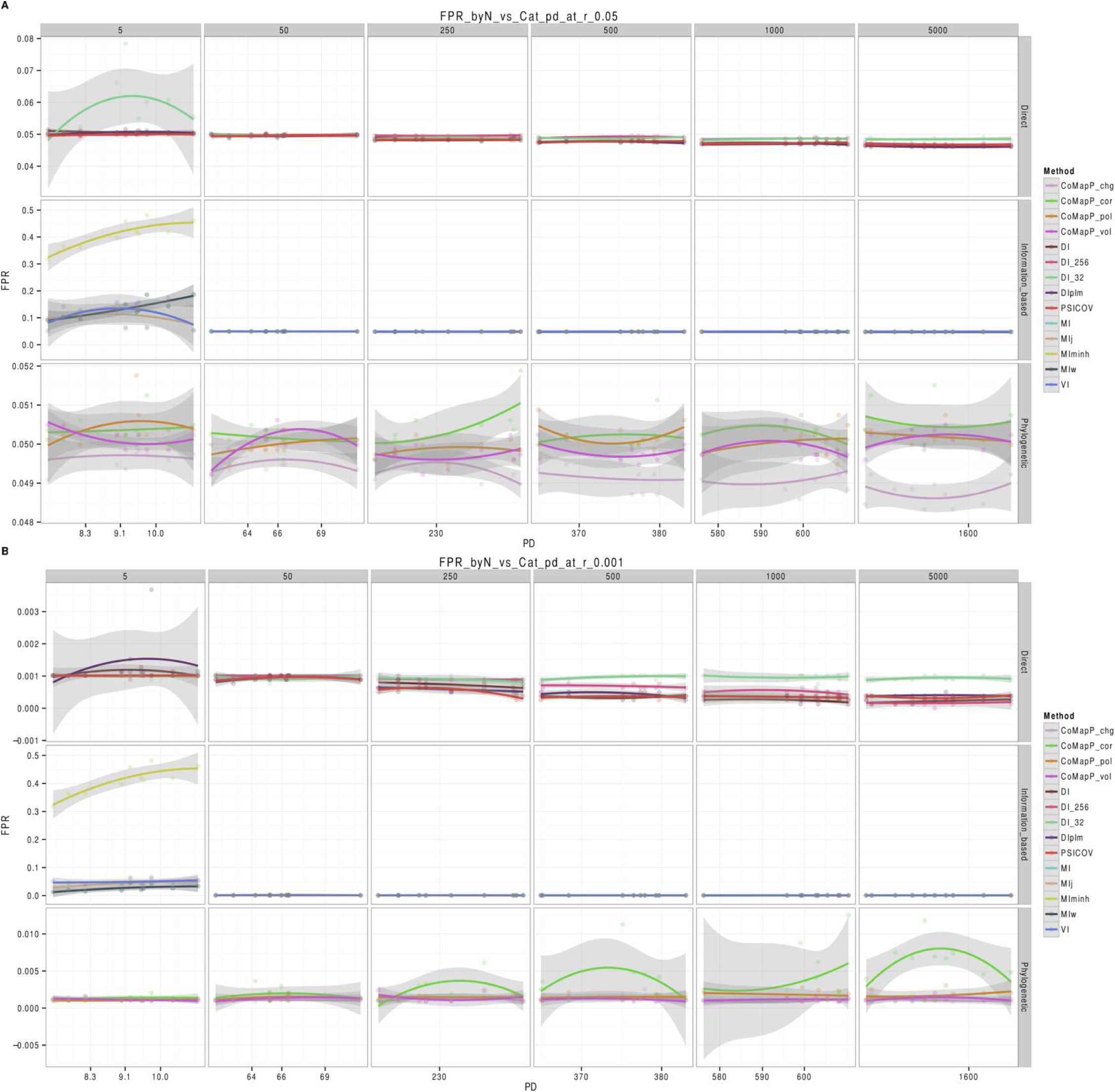
Sequence diversity may be important for controlling the false positive rate (FPR) in small alignments. **A:** Nominal FPR vs phylogenetic diversity (PD) at *P*_*empirical*_ < 0.05 and **B:** *P*_*empirical*_ < 0.001. PD is the sum of branch lengths. See Misc. Abbreviations and Table 1 for abbreviations.

**Figure S12:**
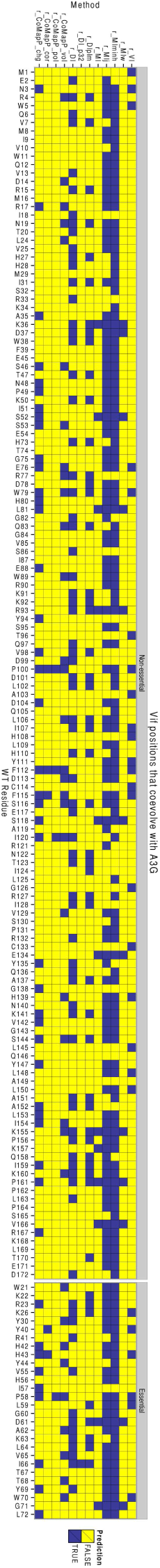
Precision (PPV) optimized predictions of contacting residues (not pairs) in Vif using previously known essential residues show varying levels of sensitivity across coevolution methods. See Misc. Abbreviations and Table 1 for abbreviations.

**Figure S13:**
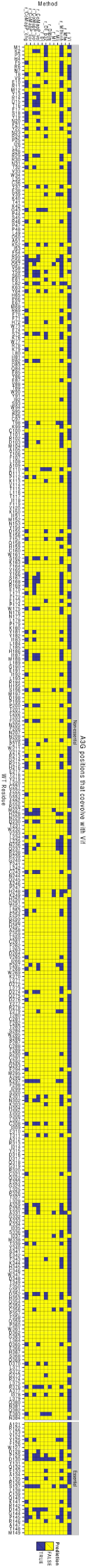
Precision (PPV) optimized predictions of contacting residues (not pairs) in A3G using previously known essential residues show varying levels of sensitivity across coevolution methods. See Misc. Abbreviations and Table 1 for abbreviations.

**Figure S14:**
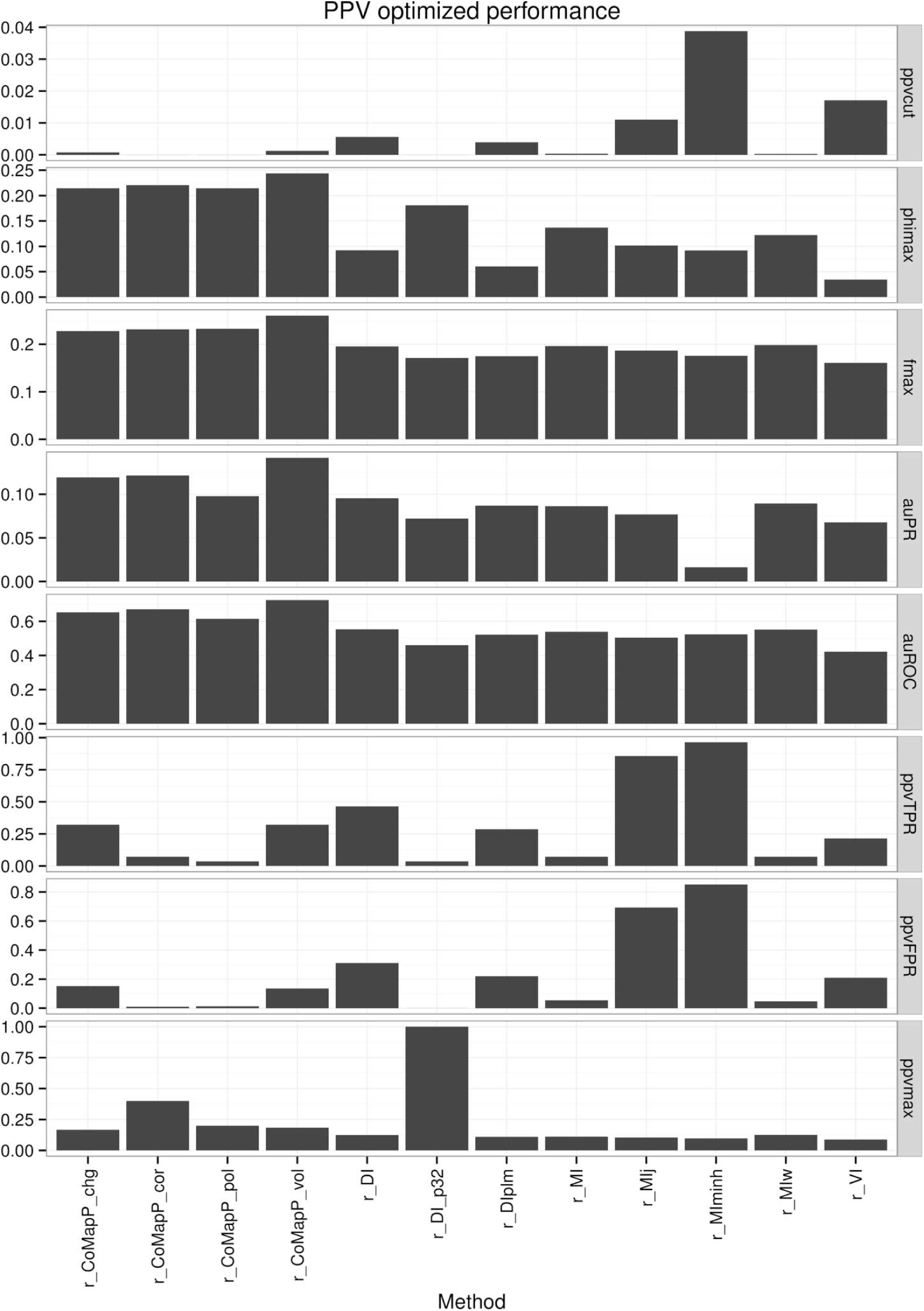
Threshold-dependent performance metrics using *P*_*empirical*_ threshold that maximizes precision in Vif. See Misc. Abbreviations and Table 1 for abbreviations.

**Figure S15:**
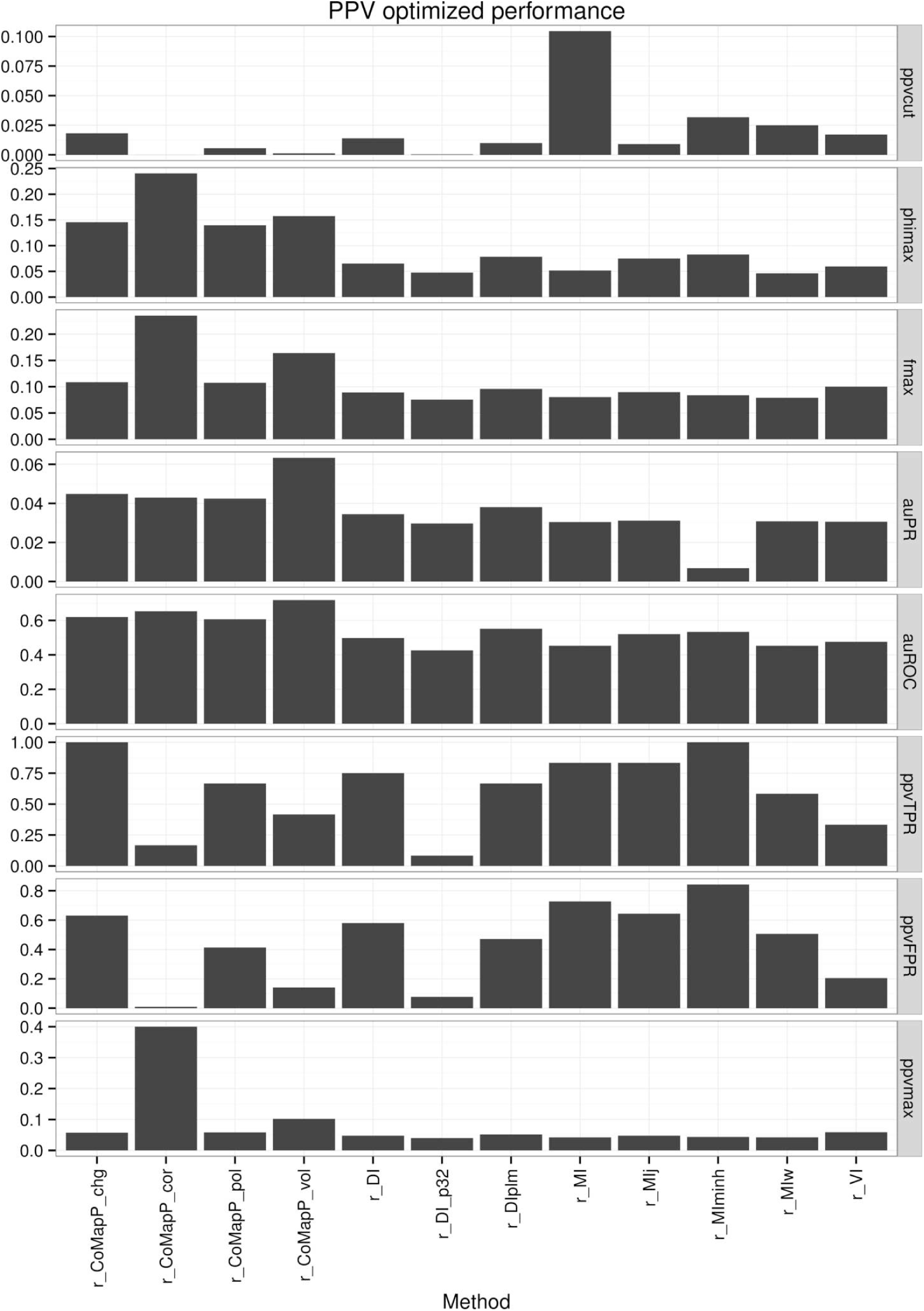
Threshold-dependent performance metrics using *P*_*empirical*_ threshold that maximizes precision in Vif using critical residues. See Misc. Abbreviations and Table 1 for abbreviations.

**Figure S16:**
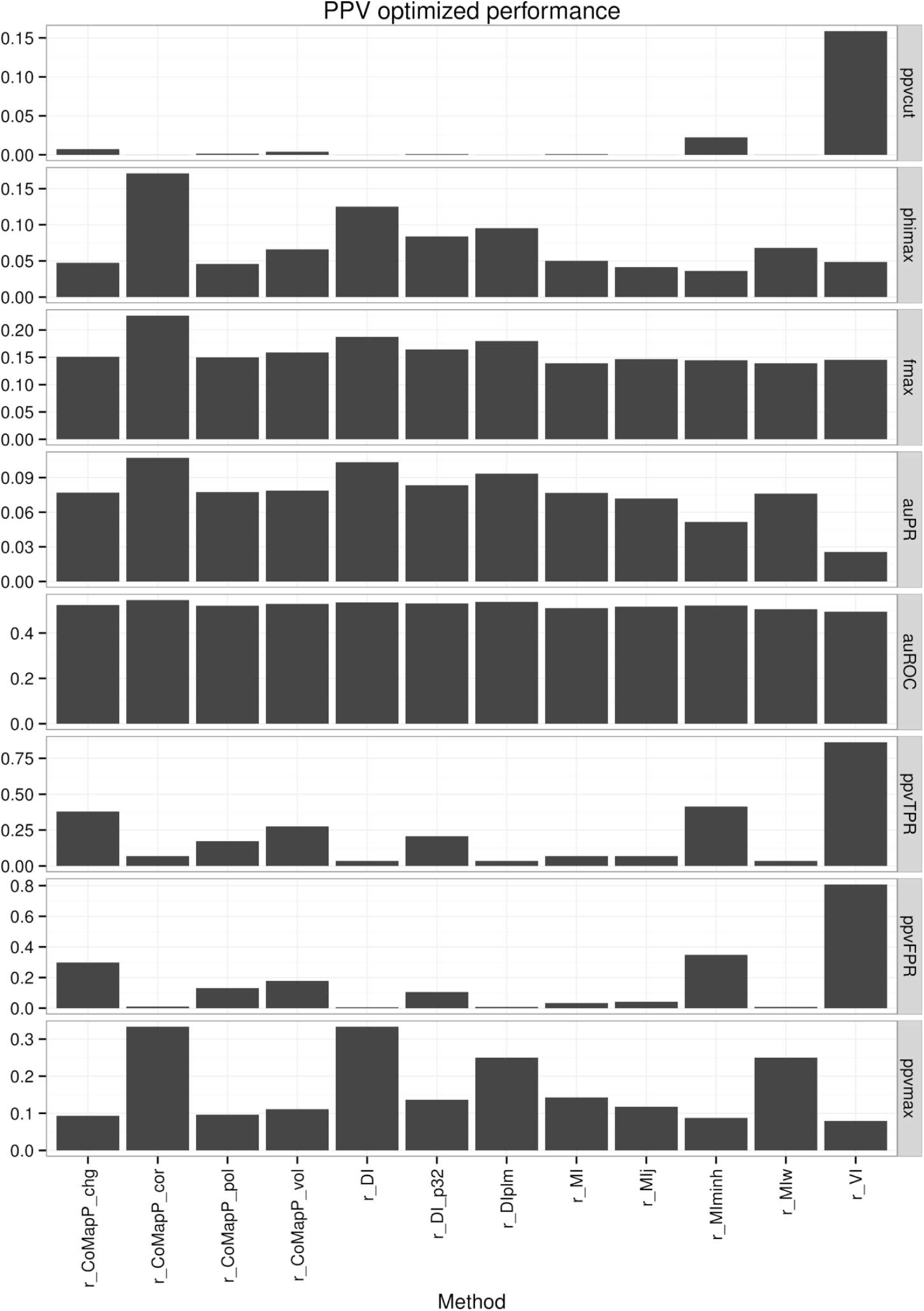
Threshold-dependent performance metrics using *P*_*empirical*_ threshold that maximizes precision in A3G. See Misc. Abbreviations and Table 1 for abbreviations.

**Figure S17:**
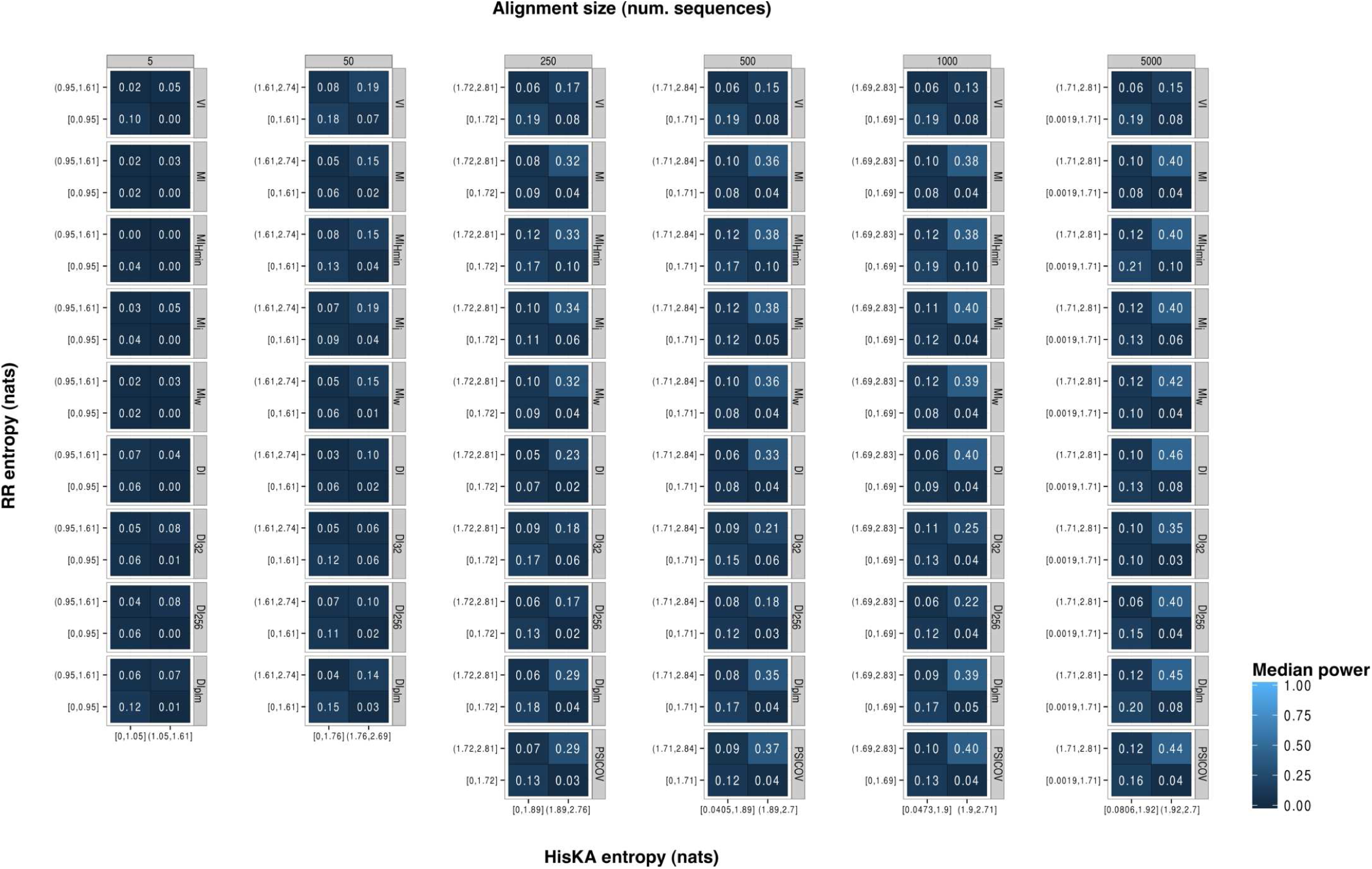
Power at FPR < 5% by HisKA-RR sub-alignment size and entropy of individual alignment columns for a subset of coevolution methods. See Misc. Abbreviations and Table 1 for abbreviations.

**Figure S18:**
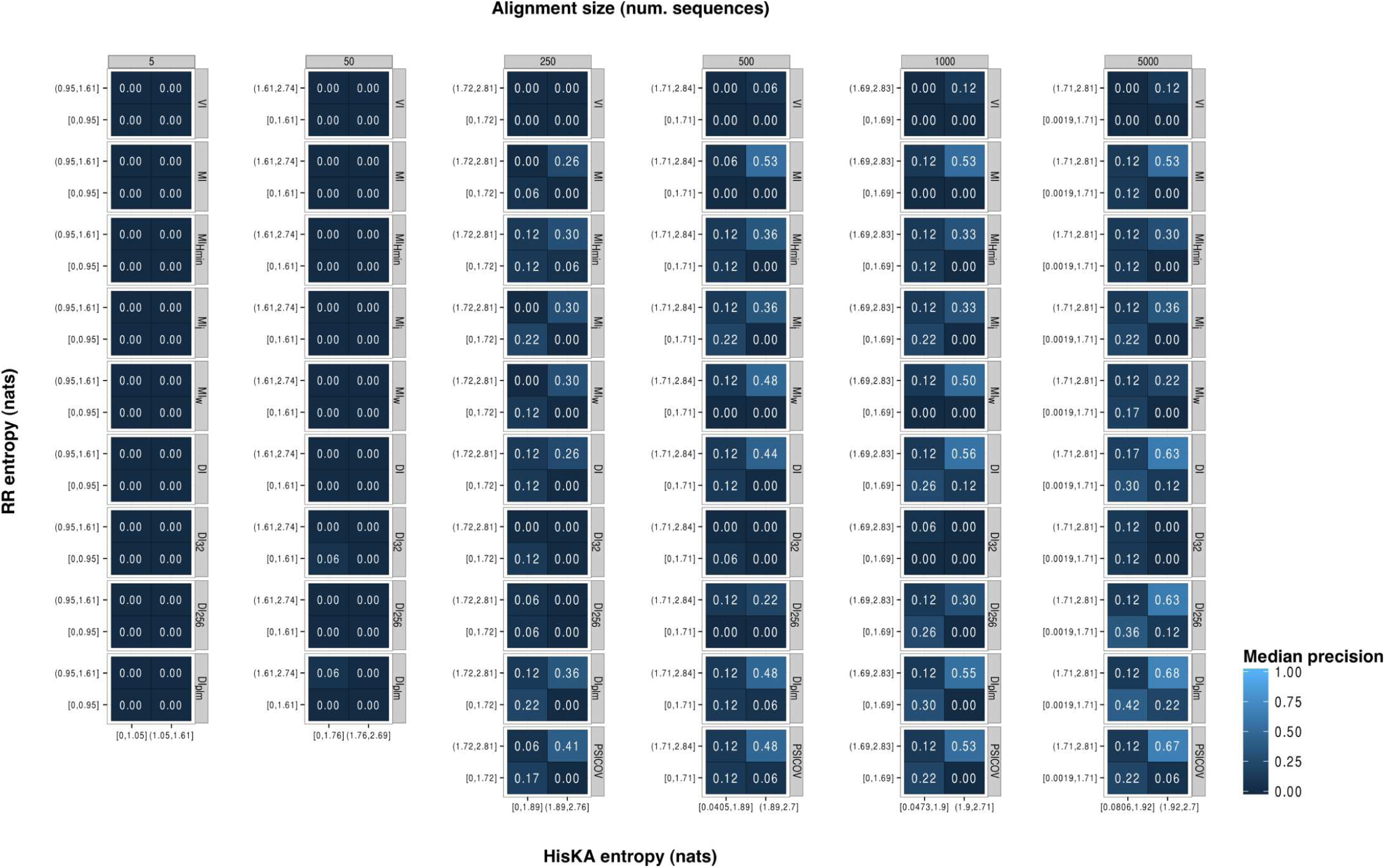
Precision at FPR < 0.1% by HisKA-RR sub-alignment size and entropy of individual alignment columns for a subset of coevolution method. See Misc. Abbreviations and Table 1 for abbreviations.

**Figure S19:**
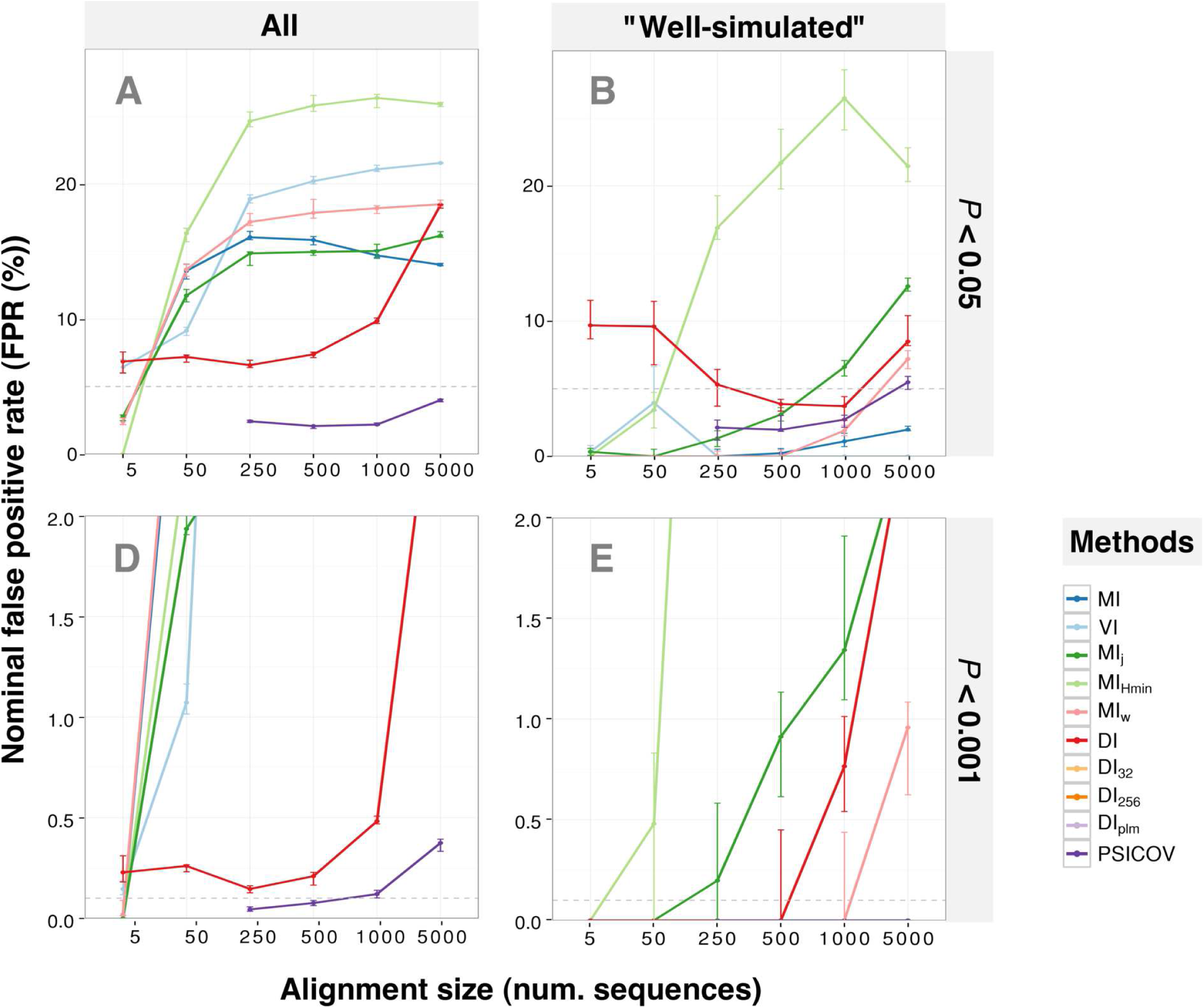
*P*_*boostrap*_ fails to control the FPR except for PSICOV at target FPR < 5% in HisKA-RR alignments. Eliminating residue pairs with large simulation errors shows PSICOV and MI_Hmin_ are most robust to variation at individual sites. See Misc. Abbreviations and Table 1 for abbreviations.

**Figure S20:**
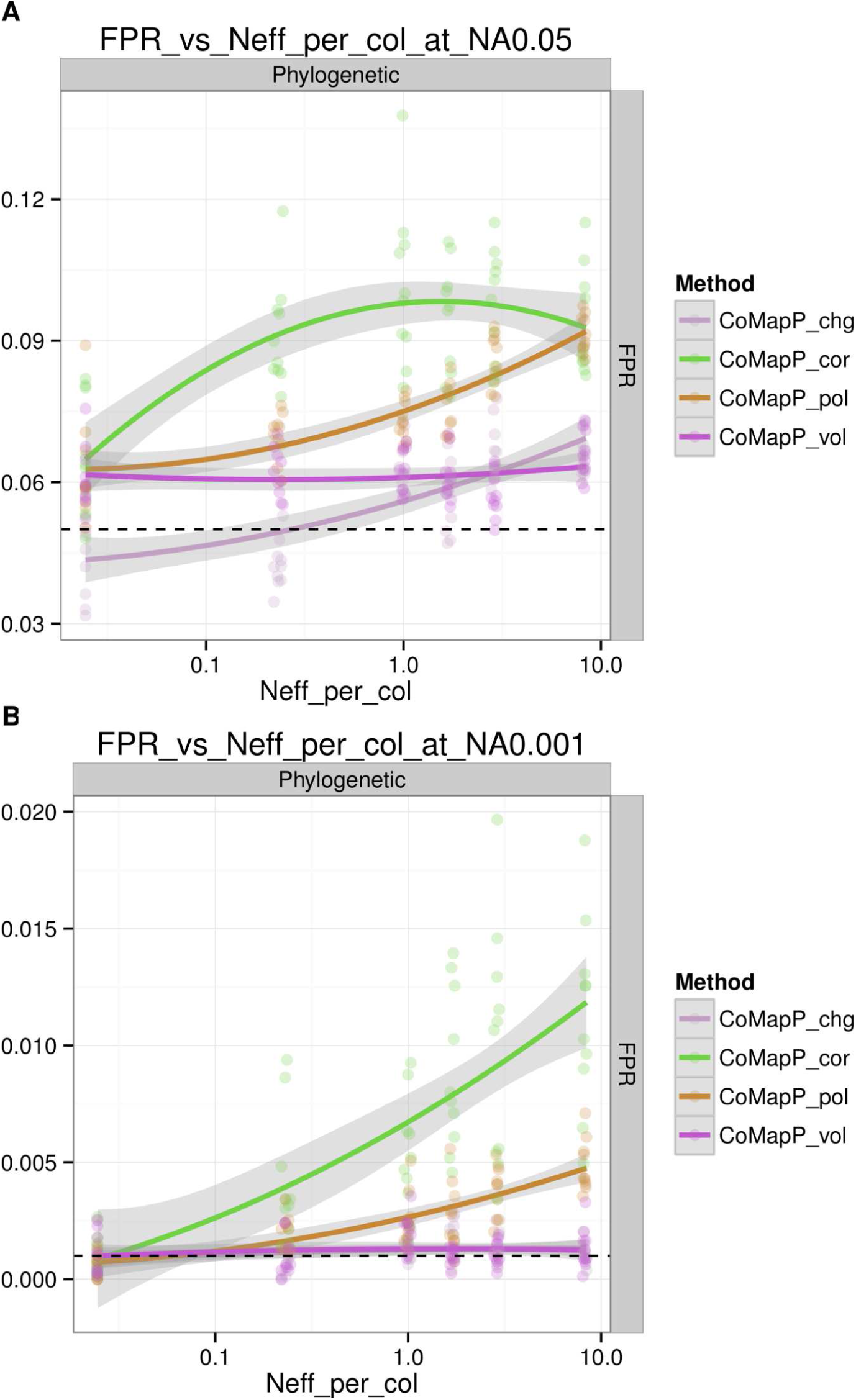
FPR vs Neff/L at 0.1% and 5% target FPRs in HisKA-RR alignments using CoMap’s internal *P*-values. See Misc. Abbreviations and Table 1 for abbreviations.

**Figure S21:**
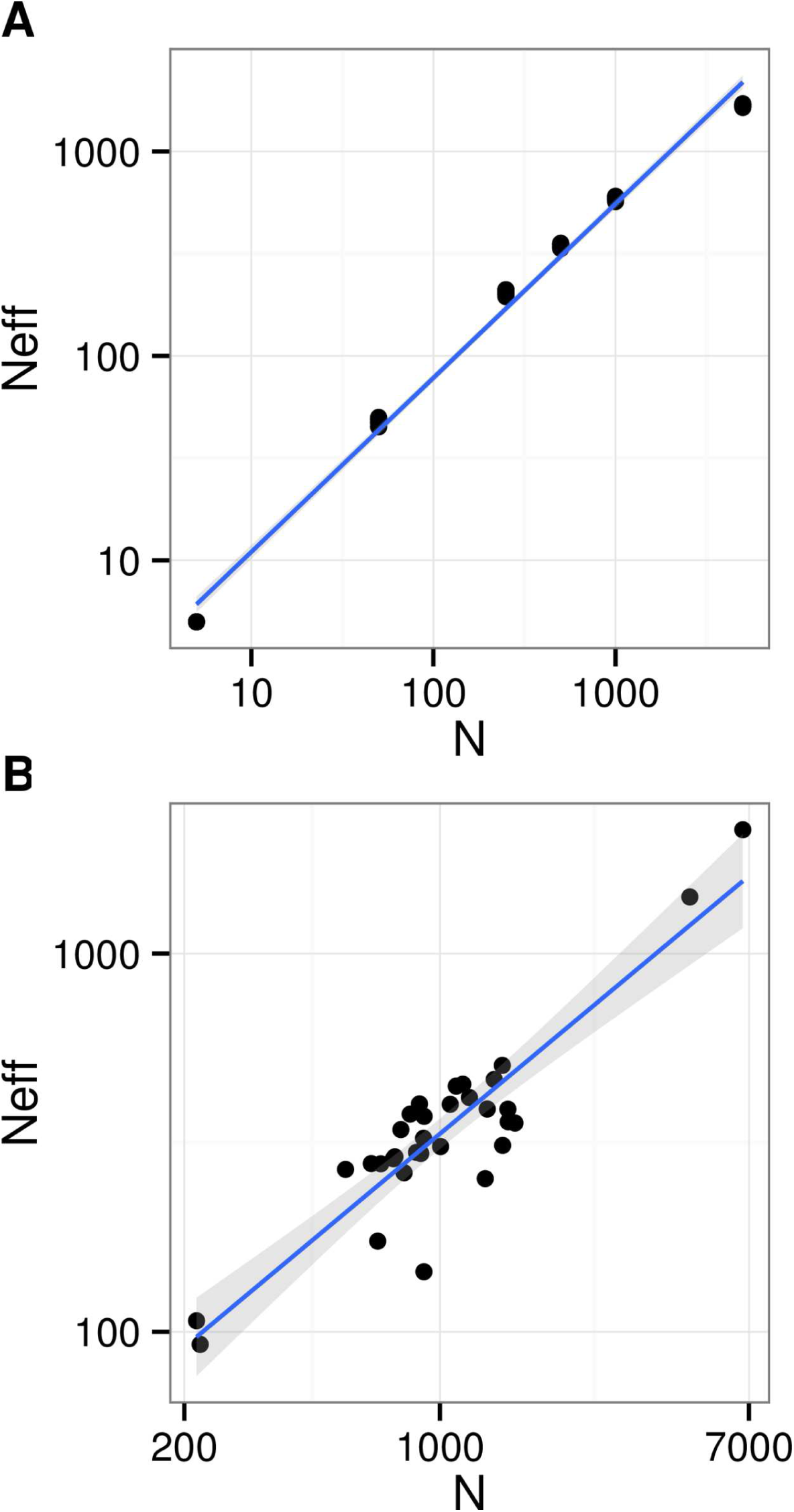
Alignment size N vs effective number of sequences as calculated by PSICOV (Neff) in **A:** HisKA-RR sub alignments and **B:** alignments in [12]. See Misc. Abbreviations and Table 1 for abbreviations.

**Figure S22:**
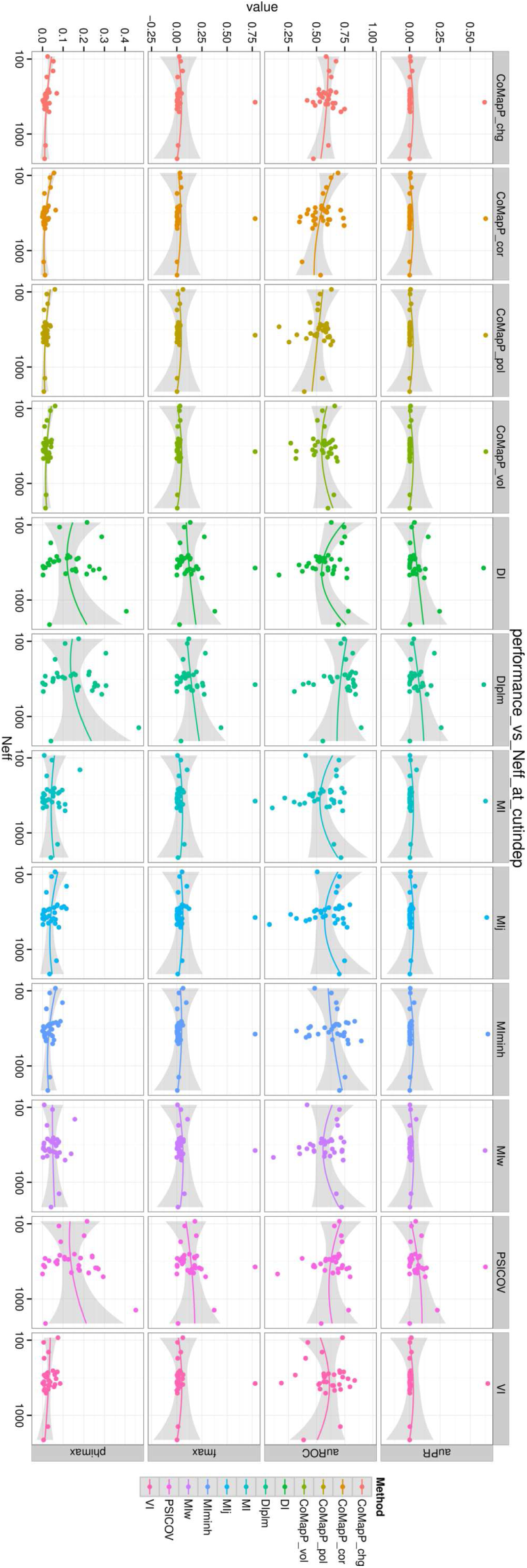
Threshold-independent performance vs Neff in alignments in [12]. See Misc. Abbreviations and Table 1 for abbreviations.

**Figure S23:**
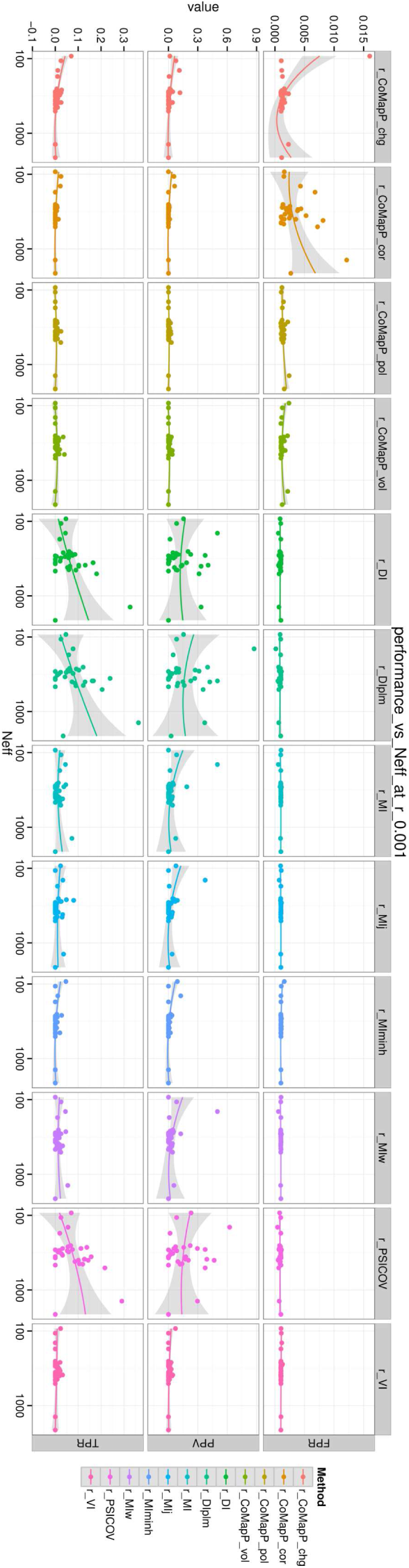
Precision (PPV) vs Neff in alignments in [12] at controlled *P*_*empirical*_ < 0.001. See Misc. Abbreviations and Table 1 for abbreviations.

**Table S1:** Coevolution method software implementation version numbers and source code.

|  | Method | Software package | Version | URL |
| --- | --- | --- | --- | --- |
| Information-based | MI | infCalc | v0.1.2 | https://github.com/aavilahe/infcalc |
|  | VI |  |  |  |
|  | MI<sub>j</sub> |  |  |  |
|  | MI<sub>Hmin</sub> |  |  |  |
|  | MI<sub>w</sub> | DCA | "2011/12" | http://dca.ucsd.edu/DCA/DCA.html |
| Direct | DI |  |  |  |
|  | DI<sub>256</sub> | Code S1 in [59] | "2013" | http://doi.org/10.1371/journal.pcbi.1003176.s002 |
|  | DI<sub>32</sub> |  |  |  |
|  | DI<sub>plm</sub> | plmDCA | symmetric_v2 | http://plmdca.csc.kth.se/ |
|  | PSICOV | PSICOV | V1.09 | http://bioinfadmin.cs.ucl.ac.uk/downloads/PSICOV/ |
| Phylogenetic | CMP<sub>cor</sub> | CoMap | 1.5.1b5 | http://home.gna.org/comap/doc/html/index.html |
|  | CMP<sub>chg</sub> |  |  |  |
|  | CMP<sub>vol</sub> |  |  |  |
|  | CMP<sub>pol</sub> |  |  |  |

**Table S2:** Species names and accession numbers of sequences used in Vif-A3G coevolution analysis.

| Mammal | A3G accession | Lentivirus | Vif accession |
| --- | --- | --- | --- |
| *Homo sapiens* | NP_068594.1 | *HIV1* | Q72499 |
|  |  | *HIV2* | Q74121 |
| *Pan troglodytes* | NP_001009001.1 | *SIVcpz* | Q1A266 |
| *Gorilla gorilla* | AAT44394.1 | *SIVgor* | ACM63194.1 |
| *Macaca mulatta* | NP_001185622.1 | *SIVmac* | P05903 |
| *Macaca nemestrina* | ADU03765.1 | *SIVmne* | AAA91932.1 |
| *Chlorocebus pygerythrus* | AEY75955.1 | *SIVver* | P27983 |
| *Chlorocebus tantalus* | AEY75957.1 | *SIVtan* | P89905 |
| *Chlorocebus sabaeus* | AEY75959.1 | *SIVsab* | AAA21506.1 |
| *Chlorocebus aethiops aethiops* | AEY75961.1 | *SIVgri* | AAA47589.1 |
| *Cercopithecus cephus* | AGE34488.1 | *SIVmus1* | ABO61046.1 |
|  |  | *SIVmus2* | ABO61055.1 |
| *Cercocebus torquatus* | AGE34491.1 | *SIVrcm* | AAK69675.1 |
| *Cercocebus atys* | AGE34496.1 | *SIVsmm* | P19506 |
| *Colobus guereza* | AGE34499.1 | *SIVcol* | AAK01034.1 |

**Table S3:**
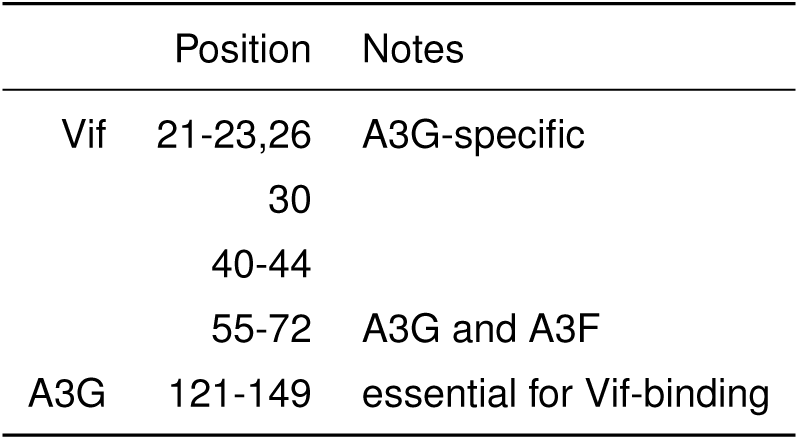
Essential and critical sites for Vif-A3G interaction.

**Table S4:** Confusion matrix definition for HisKA-RR coevolution benchmarking analysis. **True Positive Rate (TPR)**: TP / (TP + FN), **False Positive Rate (FPR)**: FP / (FP + TN), **Precision (PPV)**: TP / (TP + FP). **Phi:** (TP * TN)/sqrt((TP + FN)(TN + FP)(TP + FP)(TN + FN)), **F**: 2/(1/PPV + 1/TPR).

| $C_\beta$ distance | Prediction | |
| --- | --- | --- |
|  | Coevolving | Not coevolving |
| $< 8\text{\AA}$ | TP | FN |
| $\geq 8\text{\AA}$ | FP | TN |

**Misc. Abbreviations:** CoMap is abbreviated CMP in the main text and figures and CoMapP in supplemental figures. Effective number of sequences per column is abbreviated Neff/L. Phylogenetic distance is abbreviated PD. MI_Hmin_ appears as MIminh in figure legends. Precision (PPV) optimized metrics: ppvcut, ppvmax, ppvTPR, ppvFPR are the *P*_*empirical*_ threshold that maximizes PPV, said maximum PPV, power (TPR), and false positive rate (FPR) at said threshold.

